# Sequencing ultra-long DNA molecules with the Oxford Nanopore MinION

**DOI:** 10.1101/019281

**Authors:** John M. Urban, Jacob Bliss, Charles E. Lawrence, Susan A. Gerbi

**Affiliations:** BioMed Division (MCB Department), Brown University, Providence, RI, USA; Division of Applied Mathematics, Brown University, Providence, RI, USA; Center for Computational Molecular Biology, Brown University, Providence, RI, USA

**Keywords:** Oxford Nanopore MinION, DNA sequencing, nanopore sequencing, long reads, long DNA preparation

## Abstract

Oxford Nanopore Technologies’ nanopore sequencing device, the MinION, holds the promise of sequencing ultra-long DNA fragments >100kb. An obstacle to realizing this promise is delivering ultra-long DNA molecules to the nanopores. We present our progress in developing cost-effective ways to overcome this obstacle and our resulting MinION data, including multiple reads >100kb.

---

High-throughput DNA sequencing is at the cusp of two paradigm shifts: (1) <u>where</u> and (2) <u>how</u> sequencing is performed. The MinION from Oxford Nanopore Technologies (ONT) is a pocket-sized, long-read (>1kb) DNA sequencing device used in individual laboratories and fieldwork^1–4^, presently limited to participants in the MinION Access Programme (MAP), that detects 5mer dependent changes in ionic current, or “events” (consisting of a mean, standard deviation, start time, and duration), as single DNA molecules traverse nanopores. The events are then base-called using ONT’s Metrichor cloud service. There has been a trend of increasing base-calling accuracy and total output per flow cell^5–9^, now up to 85%^9^ and 490Mb^7^, with expectations of soon exceeding 90% accuracy and 2Gb per flow cell. Importantly, since MinION reads can be re-base-called, older reads can inherit the benefits of better base-calling.

The protocol for preparing a MinION sequencing library is still evolving, but currently includes shearing genomic DNA using a Covaris g-TUBE, an optional “PreCR” step to repair damaged DNA, end-repair, dA-tailing, adapter ligation, and His-bead purification. When properly ligated, a double-stranded (ds) DNA molecule has a Y-shaped (Y) adapter at one end and a hairpin (HP) adapter at the other. dsDNA is pulled through a pore one strand at a time starting at the 5’ end of the Y, followed by the “template” strand, and, ideally, the HP and “complement” strand. The information from either strand can be used for 1-directional (1D) base-calling, and integrating the information from both strands can be used for 2-directional (2D) base-calling, which results in higher mean quality scores (Q; Supplementary Table 1) and higher accuracy^6–11^. Molecules with a Y at each end or a nick in the template only give 1D template reads. It is also possible to obtain information from both strands but have no 2D base-calling (Supplementary Fig. 1, Supplementary Table 2). There is an approximate ratio of 1 event per base in 1D reads and 2 events per base in 2D reads (Supplementary Fig. 2). 2D reads with mean quality scores Q≥9 are considered high quality, although most other 2D reads (83-91% align to a reference^8^, ^11^) and many 1D reads (e.g., Q>3.5; Supplementary Fig. 2) are also of useful quality for various applications^2,12–17^, especially if reads are first error-corrected^7,11,16–18^ or if one does not propagate the error from base-calling by working directly with the events instead^16,18^.

The MinION has the promise to sequence ultra-long DNA fragments >100kb^19^. However, early reports suggested it falls short of this promise^5, 20^ with maximum read lengths near 10kb. In more recent reports, the majority of reads were concentrated below 30kb, but maximum read lengths were approximately 31.6kb (2D)^8^, 48.5kb (2D)^9^, 58.7kb (2D)^11^, 66.7kb (1D)^8^, 123kb (1D)^11^, and 147kb (1D)^7^. Nonetheless, reads >100kb have been rare and no instances of 2D reads >100kb have yet been reported. Here we describe modified protocols to harness the MinION’s potential for sequencing ultra-long molecules and present three resulting MinION runs (A, B, and C) with multiple reads exceeding 100kb (Fig. 1, Table 1, Supplementary Table 3).

**Figure 1:**
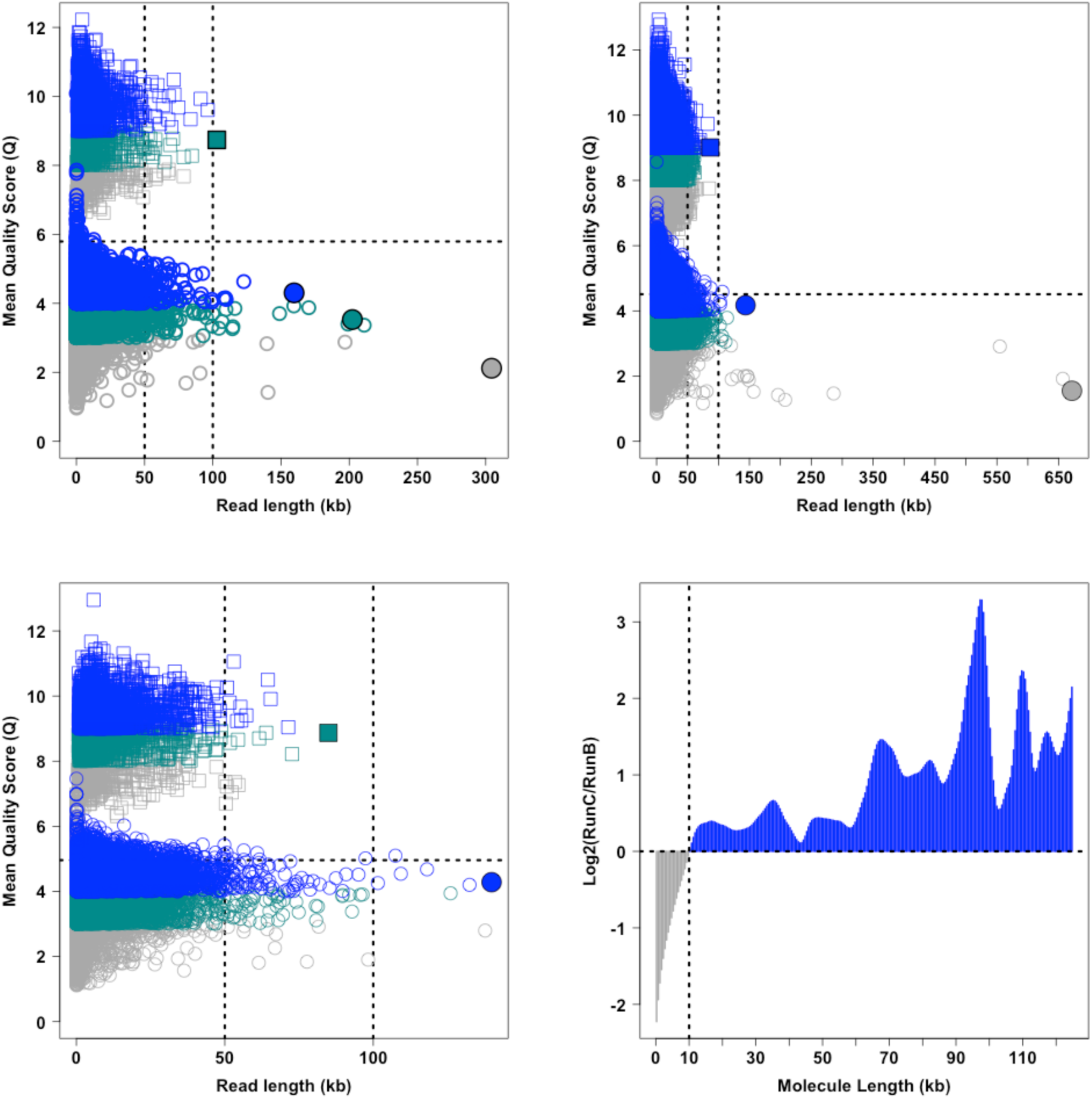
Read lengths and mean quality scores (Q) across runs. Read length vs. Mean Quality Score (Q) for **(A)** Run A, **(B)** Run B, and **(C)** Run C. In each, the distributions for both 2D reads (squares; concentrated above Q=6) and 1D reads (circles; concentrated below Q=6) are shown. For 2D reads, blue is used for reads with Q≥9, cyan for Q between 8 and 9, and grey for Q<8. For 1D reads, blue is used for Q≥4, cyan for Q between 3 and 4, and grey for Q<3. Filled-in squares and circles indicate reads highlighted in the main paper for their length and Q. The horizontal dashed line marks the minimum 2D Q. The vertical dashed lines denote 50kb and 100kb. **(D)** The log2 fold change of Run C over Run B of the proportion of total summed molecule length plotted as a function of molecule length. Run B and Run C used the same source of DNA, but differed in library preparation. Run C used a new rinse step during all AMPure clean-ups as well as took additional advantage of sequential AMPure rounds in each clean-up step. The result was depleting molecules <10kb, thereby enriching longer molecules proportionally.

**Table 1:**
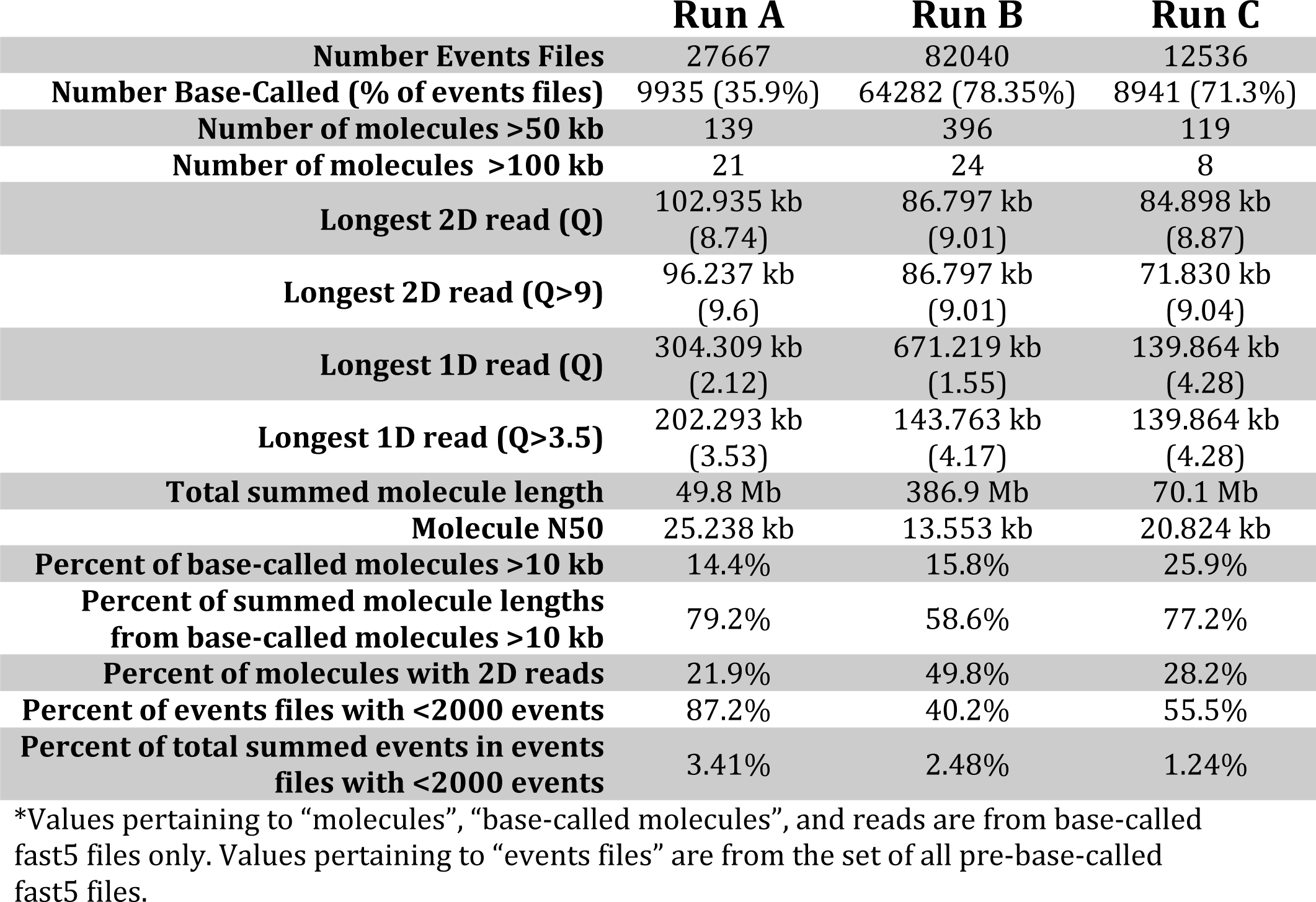
This table summarizes statistics and values discussed in the main text.

First, for Run A, we sought to maximize read lengths by gently (no vortexing) letting freshly obtained (never frozen) precipitated DNA re-suspend in TE (pH 8.0), skipping the Covaris shearing step, using wide-bored tips with gentle pipetting to minimize DNA breakage, and starting with 3X the recommended starting material to compensate for differences in molarity (Supplementary Fig. 3). Moreover, in all AMPure clean-ups we performed long elutions (20 minutes) at 37°C to help release long DNA from the beads. Run A had 9935 base-called events files featuring 139 molecules >50kb, 21 molecules >100kb, a max 2D length of 102.935kb (Q=8.74) and a max 1D length of 304.309kb (Q=2.12), although the longest 1D length with Q>3.5 was 202.293kb (Fig. 1, Table 1). The summed non-redundant molecule length was 49.8Mb with a molecule N50 of 25.238kb (Table 1). 79.2% of the summed molecule length came from molecules greater than 10kb, which made up 14.4% of all molecules sequenced (Table 1).

There were three side effects when keeping the DNA long (Table 1): (1) a low proportion of 2D reads (only ∼21.9% of base-called molecules had a 2D read); (2) a high number of tiny events files (87.2% of pre-base-called events files contained fewer than 2000 events although this made up only 3.41% of all events obtained); and (3) lower output than might have been achieved if the DNA was sheared to shorter lengths (100-400Mb routinely achieved by others in MAP at the time of our experiments). One possible explanation is that ultra-long DNA is fragile and therefore can break after end-repair leading to problems in ligation, can break after ligation leading to His-bead enrichments of HP-ligated DNA that cannot be sequenced, and can break while being injected into the MinION leading to more issues such as Y-ligated DNA that can only give 1D reads. Therefore, we proceeded to find a balance between read length, total output, and proportion of 2D reads.

For Run B we sought to minimize post-repair breaks while keeping a large proportion of the DNA >10kb (Supplementary Fig. 3). To do so we vortexed the DNA (full speed, 30 seconds) after DNA extraction (Supplementary Fig. 4A) and used normal pipette tips until end-repair, after which wide-bored tips and gentler pipetting were employed. Moreover, to compensate for the possible increase of molecules <1kb due to vortexing, two sequential 0.4x AMPure bead clean-ups were performed after end-repair. Run B had a bigger proportion of files with 2D reads (49.8%), a smaller proportion of tiny events files (40.2% of events files had <2000 events making up 2.48% of all events), and a much higher output (386.9Mb of non-redundant molecules) albeit with a lower yet still impressive molecule N50 of 13.553kb (Table 1). Run B had proportionally fewer reads >50kb than Run A, but there were more in terms of total count owing to the higher output (396 molecules >50kb, 24 >100kb). The longest 2D read was 86.8 kb (Q=9.01) and the longest 1D read was 671.219kb (Q=1.55), but the longest 1D read with Q>3.5 was 143.763kb. The majority (58.5%) of the summed molecule length came from molecules >10kb (15.8% of all base-called molecules).

For Run C, we sought to improve upon Run B by increasing the amount of data from molecules >10kb. We first explored ways to deplete DNA molecules smaller than 10kb with simple modifications to the standard AMPure bead protocol (Supplementary Fig. 4B) and found that it was sufficient to gently add a rinse after the 80% ethanol bead washes while the beads were on the magnet, incubate for 60 seconds, and remove the rinse before eluting the DNA off the magnet for 20 minutes at 37°C. We integrated rinses into the protocol for Run C in addition to doing two sequential 0.4x AMPure clean-ups before and two after end-repair (Supplementary Fig. 3) and using 4x the recommended amount of input DNA. Although Run B and Run C started with the same source of DNA, the amount of molecules <10kb in Run C was greatly depleted compared to Run B (Fig. 1D, Supplementary Fig. 5). Run C had proportionally fewer tiny events files than Run A (55.5% of events files had fewer than 2000 events) that contained proportionally fewer events (1.24%) than both Runs A and B (Table 1). There were proportionally >2-fold more base-called molecules >50kb and >100kb in Run C than in Run B, and Run C had the highest mean and median molecule sizes (Table 1, Supplementary Table 3). The longest 2D read was 84.898kb (Q=8.87) and the longest 1D read was 139.864kb (Q=4.28). The base-called molecule N50 was 20.824kb and there was a higher proportion (25.9%) of base-called molecules >10kb than both previous runs, which made up 77.2% of the summed molecule length. The percent of base-called files with 2D reads (28.2%) and the total output of the flow cell (70.1Mb of summed molecule lengths) were both intermediary between Runs A and B, suggesting that there is a trade-off between read length and output/2D reads. However, it is possible that the differences in output and proportion of 2D reads amongst the runs were due to variation in library preparations and flow cell quality, though Runs B and C had a similar estimated number of active channels (Supplementary Table 3).

Each run produced event files (69 total, Supplementary Table 5) with too many events (>1 million) for base-calling and we investigated whether they were likely to represent megabase molecules. First we eliminated all multi-million event files that contained blocks of events repeated numerous times (Supplemental Fig. 6). This error is rare (0-0.017% of all files; Supplementary Table 5), but was more prominent in files with 100 thousand to 1 million events (0-5.51%) than it was in files with <100 thousand events (0%) and was most prominent in the files with >1 million events (28.6-65.2%). Of the multi-million event files, 30 of the 69 did not contain this error (19 shown in Fig. 2A). Another concern is that large events files might arise from a faulty pore independent of a DNA molecule traversing the pore. To rule this out, we discarded 11 of the 30 remaining files that did not show evidence of a lead adapter profile (Fig. 2B). A third concern is that DNA molecules can become temporarily stuck in the pores leading to an accumulation of events from the same region of a DNA molecule. To determine how pervasive this issue might be in the remaining multi-million event files, we looked at how many times the base-caller decided two or more adjacent events corresponded to the same kmer (move=0, “stay”) instead of advancing to a new kmer (move>1) in files with <1 million events (Fig. 2C, Supplementary Fig. 7). In general, all base-called files with ≥531,779 events had an average of >80% “0 moves” in the template and complement (Fig. 2C). Indeed, the base-called sequences that have >500 thousand events appear to have come from 12.687-196.362kb molecules, also reflected in their high event:nt ratios (2-76) (Supplementary Fig. 7). Thus, it is probable that the remaining 19 files that contained 1.1-5.2 million events correspond to molecules that were stuck.

**Figure 2:**
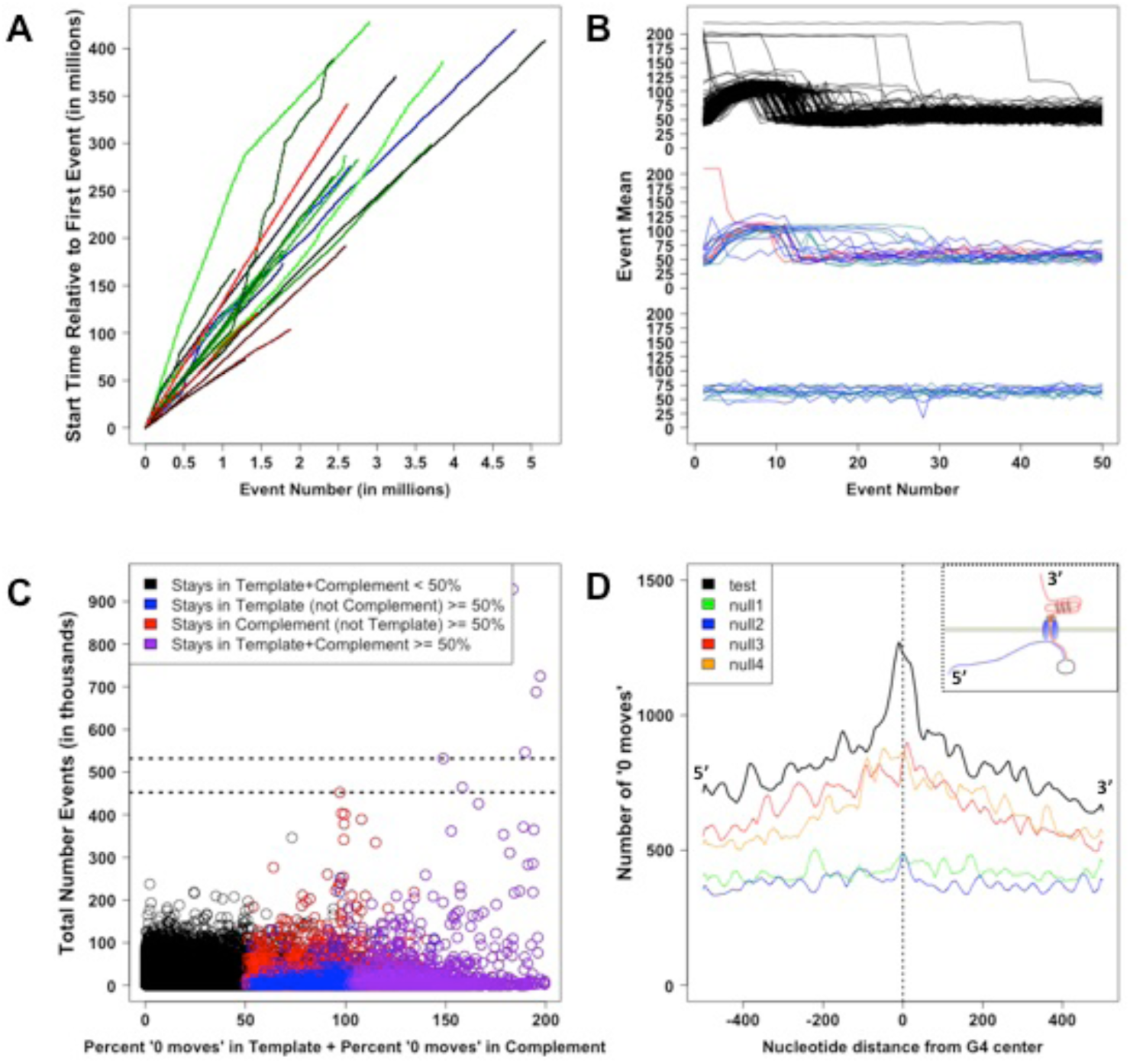
Dissecting files with >1 million events and a role of G-quadruplexes in DNA stalling. **(A)** Multi-million event files were first filtered to remove files with artificially high event counts due to repeated blocks of events. Shown are the start times as a function of event number for the multi-million event files that did not have repeated blocks of events and that had evidence of the lead adapter (i.e. middle row of 2B). Different colors represent different molecules. Shades of blue are from Run A, shades of green are from Run B, and shades of red are from Run C. **(B)** The event means of the first 50 events for: (Top) 150 randomly sampled (50 from each run), high quality (Q≥9) 2D reads; (Middle) multi-million event files that did not have repeated events and do show evidence of the lead adapter; (Bottom) multi-million event files that did not have repeated events and do NOT show evidence of the lead adapter. Red lines are from Run A, blue lines are from Run B, and cyan lines are from Run C. **(C)** The total number of events plotted as a function of the summed percentages of base-called events in the template and complement (T and C) reads that had “0 moves”. Notably all files with ≥452301 events (bottom dashed line) had on average ≥50% base-called events with “0 moves” in the T and C reads and all files with ≥531779 events (top dashed line) had on average ≥80% base-called events with “0 moves” in the T and C reads. **(D)** The number of “0 moves” as a function of proximity to G4 motif centers (test condition) or the centers of randomly selected positions (null 1-4 conditions). Briefly, positions were randomly selected on any template or complement (T or C) read without a G4 motif (‘null 1’), on any T or C read including those with G4 motifs (‘null 2’), on the same read the G4 motif originated (‘null 3’), and on the same read the G4 motif originated ensuring it did not overlap the G4 motif (‘null 4’). The inset at the top right depicts the orientation (5’ – 3’) of a DNA molecule going through a pore from the top chamber to the bottom chamber and how a G4 might cause a DNA molecule to stall.

Finally, we sought to understand how DNA might get stuck in pores. We hypothesized one possibility is that a highly stable DNA secondary structure known as a G-quadruplex (G4)^21^ may form and block further translocation until it unfolds (Fig. 2D, Supplementary Fig. 8), resulting in an accumulation of measurements of a 5mer or set of 5mers slightly upstream of the G4. Indeed, there is a significantly higher number of “0 moves” near G4 motifs than near randomly selected locations even when controlling for read-specific effects (Supplementary Table 6A). Moreover, there is a significantly higher number of “0 moves” near G4 motifs on complement strands than near G4 motifs on template strands consistent with the higher propensity of G4-folding in single-stranded DNA^21^ (Supplementary Table 6B). With each additional poly-G tract inside a G4 motif there are additional ways a G4 can form, increasing the probability that one will. Thus, if G4 structures are associated with DNA stalling, one might expect more “0 moves” near G4 motifs with more poly-G tracts. Indeed, there is also a significantly higher number of “0 moves” near G4 motifs that have >4 poly-G tracts than G4 motifs with only 4 G-tracts (Supplementary Table 6C). Finally, in an aggregate analysis looking at all template and complement reads, there is a clear enrichment of “0 moves” near G4 motifs with the highest enrichment (Run A) and shoulders (Runs B, C) slightly upstream (-9 to -27 nt) of the G4 motif (position 0) as expected (Fig. 2D, Supplementary Fig. 8). Nonetheless, it is clear that G4s are not the only way DNA molecules get stuck since there are many “0 moves” far away from and on different molecules than G4 motifs. Fortunately, since the base-caller can identify “0 moves”, it is able to deal with DNA stalling, though there is a slight decrease in Q with each increase in the proportion of called events that have “0 moves” (Supplementary Fig. 7). Consistently, reads with G4 motifs appear to have a slightly lower average Q than all reads (Supplementary Table 6D). Nonetheless, in future studies, “0 moves” might serve as an indicator of whether and how often all possible G4 motifs in a genome form *in vitro* in a single MinION experiment.

In conclusion, our data demonstrate that with our modified protocols the MinION can sequence as many ultra-long DNA molecules >100kb that make it intact to the nanopores. Importantly, we demonstrate that it is possible to obtain high quality 2D reads >100kb (e.g. 102.935kb, Q=8.74). Indeed, using our modified protocols, ONT internally obtained a 192kb high quality 2D *E. coli* read that mapped to the reference genome. The modified protocols presented here will help others obtain similar read size distributions.

## ACKNOWLEDGMENTS

We thank Oxford Nanopore Technologies for including us in MAP, the MAP community for online discussions, Ben Raphael for providing computing resources for the MinION, and Mark Howison for helpful discussions. We gratefully acknowledge support from predoctoral fellowships from NSF GRFP (DGE-1058262) and NSF EPSCoR (grant# 1004057) to J.M.U.

## AUTHOR CONTRIBUTIONS

J.M.U. conceived, designed, and carried out the experiments, analyzed the data, wrote and utilized the associated “poreminion” software (https://github.com/JohnUrban/poreminion), and prepared the manuscript. J.B. performed and supplied mass-matings of flies for embryo collection and contributed to discussions about the project. C.E.L. provided guidance in statistics and experimental design. S.A.G. provided guidance in molecular techniques and experimental design, and helped prepare the manuscript. All authors read and approved the manuscript.

## COMPETING FINANCIAL INTERESTS

JMU and SAG are members of the MinION Access Programme (MAP) and have received free reagents for nanopore sequencing.

## Supplementary Materials

### Supplementary Methods

#### Overview of genomic DNA isolation, MinION sequencing, and analysis

We have been developing our protocols and reference-free analyses while collecting MinION data for a future assembly of the fungus gnat (*Sciara coprophila*) genome. All libraries were prepared with the SQK-MAP004 kit and reads were base-called with Metrichor 1.12 r7.X 2D-basecalling XL. In our analyses, we characterize the full distribution of reads as well as the pre-base-called events files. Reads were filtered to remove those with errors in event timing (repeated blocks of events) and those that were not base-called due to having too few events (<200), too many events (>1 million), or “no template” (Supplementary Table 5). The rare error that leads to blocks of events being repeated numerous times is easily identified by events with earlier start times than preceding events and therefore we often refer to it as the “time error”. It is important to point out that the software issue that led to this error in a small number of files has since been resolved and will not occur in future experiments. Filtering and data analyses were carried out using our open source MinION toolset called “poreminion” (https://github.com/JohnUrban/poreminion) as well as in R^1^ and using poretools^2^.

#### DNA Extraction

Genomic DNA (gDNA) was extracted from 51 mg (sample 1, Run A) and 53 mg (sample 2, Runs B and C) of synchronized 2 day old *Sciara* male embryos from ∼30 mass matings each that were extensively washed with sterile TE, pH 8.5. There were 5-8 adult males and 10 male-producing adult females per mass mating (note that Sciara females have either only male or only female offspring, which we can control). The washed male embryos were homogenized with a blue pestle inside a 1.5 ml microfuge tube in 200 μl DNAzol (Life Technologies): ∼5-10 gentle strokes for sample 1; ∼20 strokes for sample 2. Then 800 μl more DNAzol was added to the tube for 1 ml DNAzol total. The homogenate from sample 2 (not sample 1) was vortexed (full speed, 30 seconds) to further facilitate homogenization and lysis. The DNAzol homogenate was (for both samples 1 and 2) incubated with 5 μl RNase A Solution (Qiagen) for 10 minutes at 37°C, followed by 5 μl Puregene Proteinase K (Qiagen) for 10 minutes at 37°C. The DNAzol homogenate was then centrifuged at 10,000xg for 10 minutes to pellet debris. The supernatant was transferred to a new 1.5 ml tube [using a wide-bored tip (Axygen: Max Recovery, DNase/RNase-free, Sterile, Wide-bore tips) for sample 1 and regular tip (Olympus: Low-binding, RNase/DNase-free, Sterile, Barrier tips) for sample 2] and 500 μl 100% ethanol was added. The tube was capped and slowly inverted 50 times, incubated at room temperature for 2 minutes, then on ice for 2 minutes. Precipitated DNA was pelleted at 18,000xg for 10 minutes. The DNA pellet was washed twice with 80% ethanol, air-dried for 30 seconds, and re-suspended in 1xTE, pH 8. Sample 2 (not sample 1) was vortexed (full speed, 30 seconds) to help facilitate re-suspension. Furthermore, to help re-suspension of high molecular weight DNA, both samples were incubated at 37°C for 1 hour before an overnight incubation at room temperature and storage at 4°C (in 1X TE, pH 8) for up to 2 weeks before use.

#### AMPure beads clean-up #1

After DNA extraction, but before beginning the MinION library preparation, an appropriate volume of the DNAzol-extracted DNA was used to obtain 3, 3.6, and 4 μg of gDNA for Runs A, B, and C respectively. For all runs (A, B, and C), the DNA was cleaned with 1.0x AMPure beads (Beckman Coulter Agencourt) with the following specifications. An equal volume (1.0x) of AMPure beads was added to DNA in 1X TE (pH 8), incubated for 15 minutes at room temperature (RT), then pelleted on a magnetic rack for 5 minutes (RT) before removing the supernatant. The beads were then washed with 500 μl buffered 80% ethanol (always buffered with 10 mM Tris-Cl, pH 8) followed by a second wash with 200 μl buffered 80% ethanol, a brief gentle spin in a microfuge to collect any remaining 80% ethanol at the bottom of the tube, 1 minute re-pelleting on a magnet, removal of the remaining buffered 80% ethanol collected at the bottom of the tube without disturbing beads, air drying 7 minutes, re-suspending in 175 μl Ultra Pure Water (UPW, Life Technologies UltraPure DNase/RNase-Free Distilled Water) at 37°C for 20 minutes to help ensure long DNA elutes off the beads, pelleting the beads on a magnet for 5 minutes, then transferring 174 μl supernatant with DNA to a new tube (for PreCR). For Run A, wide-bored tips and gentle pipetting were used. For Runs B and C, normal tips and normal pipetting were used. For Run C, directly before eluting the DNA off the beads, a “rinse” of 200 μl 10 mM Tris (pH 8) was gently added to the tube wall opposite the magnetically pelleted beads, incubated at room temperature for 60 seconds, and gently removed. This is an additional rinse step used to help deplete DNA <10kb (see Supplementary Fig. 4B, Fig. 1D, and Supplementary Fig. 5).

#### PreCR DNA Repair

Since 3-4x the ONT-recommended amount of starting material was used, PreCR (New England BioLabs, NEB) was performed in double the volume, with double the reagents, for double the time (all relative to ONT protocol): 200 μl total volume with 174 μl AMPure cleaned DNA, 20 μl 10x Thermopol buffer, 2 μl 100x NAD+, 2 μl 10 mM dNTPs, and 2 μl PreCR Repair Mix. The reaction was incubated at 37°C for ≥60 minutes.

#### AMPure beads clean-up #2

For Runs A and B, we proceeded as in clean-up #1, only with a 0.4x AMPure beads ratio and transferring 85 μl to a new tube at the end (for End-Repair). For Run C, sequential 0.4x clean-ups were performed in the following way. In the first 0.4x AMPure clean-up, the beads were air-dried for only 2 minutes (after the buffered 80% ethanol washes) before adding 140 μl 0.4x AMPure solution (100 μl UPW, 40 μl AMPure beads), gently re-suspending the beads in the second 0.4x solution, and proceeding as in clean-up #1 for Run C. Importantly, a “rinse” was again performed before eluting DNA off the beads. Specifically, 100 μl 10 mM Tris-Cl (pH 8) was added to the tube wall opposite the pelleted beads, and incubated for 30 seconds at room temperature while the tube remained on the magnet before gently removing the rinse. Again (for all runs), elution off the beads (into 85 μl UPW) was performed for a longer time and a higher temperature than manufacturer recommendations to facilitate the elution of long DNA (≥20 minutes, 37°C).

#### End-Repair

The NEBNext End-Repair Module (NEB) was used: 85 μl of DNA from the previous AMPure step, 10 μl NEBNext End-Repair Reaction Buffer (10X), and 5 μl NEBNext End-Repair Enzyme Mix. The reaction was incubated for 30 minutes at 22°C.

##### AMPure beads clean-up #3

For Run A, we proceeded as in #2, except that elution was in 30 μl. For Run B, two sequential 0.4x AMPure clean-ups were performed as done for Run C in #2 (but with no rinse). For Run C, we proceeded as Run C in #2 with 2 sequential 0.4x washes and the rinse step. Specifically, 100 μl 10 mM Tris-Cl (pH 8) was added to the tube wall opposite the pelleted beads, then incubated for 30 seconds at room temperature while the tube remained on the magnet before gently removing the rinse and eluting for ≥ 20 minutes at 37°C. For all runs, DNA was eluted into 30 μl 10 mM Tris-Cl (pH 8).

##### dA-tailing

The NEBNext dA-Tailing Module (NEB) was used: 25 μl DNA from the previous AMPure step, 3 μl NEBNext dA-Tailing Reaction Buffer (10X), and 2 μl Klenow Fragment (3´ → 5´ exo–). The reaction was incubated for 30 minutes at 37°C.

##### Adapter Ligation

The following were combined: 30 μl dA-tail reaction, 8 μl UPW, 10 μl ONT SQK-MAP004 Adapter Mix, 2 μl ONT SQK-MAP004 HP adapter, 50 μl 2X Blunt/TA Ligase Master Mix. The reaction was incubated for 30 minutes at 22°C.

##### Enrichment of HP-ligated DNA with His-Beads

550 μL UPW was added to 550 μL SQK-MAP004 2X Wash Buffer (this is called “1X Wash Buffer”), then mixed by inverting 10 times, and briefly spun down in a microfuge. “His-beads” (Dynabeads His-tag Isolation and Pulldown; Life Technologies) were re-suspended by vortexing for 30 seconds. Then 10 μl of re-suspended beads was transferred to a 1.5 ml Protein LoBind tube (Eppendorf), and combined with 250 μl 1X Wash Buffer. The tube was placed on a magnet for 2 minutes before aspirating off the supernatant. The his-beads were re-suspended in another 250 μl 1X Wash Buffer and placed on the magnet for another 2 minutes before removing the supernatant. The twice-washed and pelleted his-beads were re-suspended in 100 μl undiluted (2X) Wash Buffer and are referred to as the “washed his-beads”. 100 μl “washed his-beads” was added to the 100 μl adapter ligation reaction, mixed by gentle pipetting (wide-bore tips), and incubated at 22°C for 5 minutes. The his-beads were then pelleted using a magnetic rack for 2 minutes before removing the supernatant, then washed twice with 250 μl 1X Wash Buffer (with 30 second incubations). The tube was briefly spun in a microfuge to collect excess Wash Buffer at the bottom of the tube, which was then placed in the magnetic rack for 2 minutes before removing the excess buffer. The pelleted his-beads were re-suspended in 30 μl of Elution Buffer with gentle pipetting using a wide-bored tip, adding the buffer close to the his-beads (to avoid any residual wash buffer on the sides of the tube). The elution was incubated for 10 minutes at 22°C before pelleting with a magnetic rack for 2 minutes. The eluted supernatant (called the “Pre-Sequencing Mix” (PSM)) was transferred to a new Protein LoBind tube.

##### Pre-sequencing Mix (PSM)

Before the PreCR step, Run A started with 3000 ng gDNA, Run B started with 3600 ng, and Run C started with 4080 ng. All had A[260/280] and A[260/230] >1.8 prior to PreCR and >2.0 after subsequent clean-up steps during the library preparation as determined by Nanodrop (Thermo Scientific). Each sample preparation ended with between 300-400 ng DNA in the PSM as measured by Qubit dsDNA HS (Life Technologies).

##### Loading the Sequencing Mix (SM)

Sequencing Mix (SM) was made using 3-6 μl PSM, 3-4 μl Fuel, and EP buffer up to 150 μl. SM was made fresh for loading/re-loading at various intervals 6-7 times throughout each run. See Supplementary Table 4B for exact details.

##### Base-calling

MinION events data for each DNA molecule is stored in an individual “fast5” file. All fast5 files were base-called using the Metrichor 1.12 r7.X 2D-basecalling XL protocol. Metrichor returns fast5 files that are updated with base-calling information.

##### Filtering base-called fast5 files

Metrichor (ONT base-caller) returns updated fast5 files into two folders: “pass” and “fail”. “Pass” contains only fast5 files where 2D base-calling was successful and the mean quality of the 2D read is ≥9. Everything else (including other fast5s containing 2D reads with Q<9, fast5s with only 1D base-calling, and fast5s that failed base-calling) goes into the “fail” folder. To analyze all base-called molecules, we filtered the contents of the fail folder to remove un-basecalled files and further filtered to remove any base-called files that contained the “time error” where a block of events is repeated. Both were accomplished by using our toolset for working with MinION data, called “poreminion” (which, for some of its functionality, sources the Fast5File classes from poretools^2^):

$ poreminion uncalled -m -o fail-filter fail/

$ poreminion timetest -m -o fail-filter fail/

The syntax for both subcommands is:

poreminion subcommand options (-m –o outprefix) target-directory (to search).

The “uncalled” subcommand identifies all fast5s that were not base-called due to either: (1) too many events, (2) too few events, or (3) no template found. The flag “-o” gives a prefix to poreminion, which writes text files containing the names of the fast5s in each of the above categories in addition to a summary statistics file describing how many fast5s were searched, how many were assigned to each category, as well as the minimum, maximum, median, and mean number of events found in files of each category. It also reports the number of events for all files with too many events to base-call. The “-m” flag tells poreminion to not only report the names of the un-basecalled files, but to also move them into their own folders. The “timetest” subcommand searches for files with repeated blocks of events, which are identified by looking at the start times of all events in a fast5 file. If a fast5 file contains an event start time that is earlier than the event that preceded it, then the fast5 file contains this error and is reported (“-o”) and moved to a folder for files with this error (“-m”).

##### Obtaining molecule size, mean quality score (Q), other statistics, and plotting

Each base-called fast5 file from a MinION run describes data from a single molecule, yet there can be up to 3 reads per file: template, complement, and 2D. We define the molecule size as the length of the 2D read if present, the length of the template read if there is only a template read present, and the length of longer of template and complement reads when both are present in the absence of a 2D read. Thus, the only time there is a choice is in the latter situation. The majority (68-86%, Supplementary Table 2, Supplementary Fig. 1) of files with a template and complement sequence have a 2D read, all of which have a template:complement sequence length ratio between 0.5 and 2. Moreover, most files that contain both template and complement sequences with a sequence length ratio between 0.5 and 2 have 2D reads (Supplementary Fig. 1). This means that when a choice needs to be made between the template and complement they are vastly different sizes. The complement can be much smaller than the template, for example, when there is a nick in the complement strand. The template can be much shorter than the complement, for example, when the motor protein on the Y-adapter falls off allowing the template to zip through until it is caught by the hairpin motor protein. In these situations, the longer read better represents the size of the molecule that was sequenced. Importantly, molecule size allows for a single/non-redundant length from each base-called fast5 file (i.e. single molecule) to compute statistics on, such as the summed length of sequenced molecules and molecule N50, in addition to statistics computed on all read types. This molecule size estimate as well as many descriptive metrics of the fast5 file were obtained with the poreminion subcommand “fragstats” after combining all base-called files from the pass and fail folders.

$ poreminion fragstats all-basecalled-files/ > fragstats.txt

The “fragstats” subcommand gives a tab-delimited output where each line describes an individual fast5 file (or single molecule) with the following columns (for version 0.4.3).

1 = read name

2 = estimated molecule/fragment size

3 = number input events

4 = if complement detected

5 = if 2D detected

6 = number of template events

7 = number of complement events

8 = length of 2D sequence

9 = length of template sequence

10 = length of complement sequence

11 = mean quality score of 2D sequence

12 = mean quality score of template sequence

13 = mean quality score of complement

14 = ratio of number template events to number complement events

15 = channel number molecule traversed

16 = heat sink temperature while molecule traversed

17 = number of called template events (after events pruned during base-calling)

18 = number of called complement events (after events pruned during base-calling)

19 = number of skips in template (number 0)

20 = number of skips in complement (number 0 moves)

21 = number of stays in template

22 = number of stays in complement

23 = strand score template

24 = strand score complement

25 = number of stutters in template

26 = number of stutters in complement

The tab-delimited fragstats.txt file was then brought into R to make most plots (Fig. 1, Supplementary Figures 1, 2, 3, and 7). The fragstats file was also summarized (generating many of the statistics reported such as in Table 1 and Supplementary Tables 1A-C, 2, 3A-E) using:

$ poreminion fragsummary –f fragstats.txt

##### Analyzing files with too many events (>1 million) to base-call

###### Time Error

Poreminion was used to extract the event start times from the multi-million event fast5 files into text files (columns of poreminion ouput = event mean, event standard deviation, event start time, event duration), which were brought into R^1^ for visualization.

$ poreminion events –f5 target.fast5 | cut –f 3 > start.times.txt

In the directory with the fast5 files that had too many events to base-call (this directory was made by “uncalled” filtering above), the poreminion “timetest” subcommand was used to further filter these multi-million event fast5s to keep only those without the ‘time error’. Time errors (and lack thereof) were visualized in R from events text files extracted with the above poreminion command.

###### Lead adapter

The first 50 events of the remaining multi-million event fast5 files were obtained with poreminion (poreminion events –f5 target.f5) and searched for evidence of the lead adapter event mean profile by comparing to 150 randomly sampled (50 from each run A,B,C) pass fast5s (containing high quality 2D reads). Since there were so few multi-million event files, it was sufficient to manually separate ones with the lead adapter profile from ones that did not. However, we also found that the simple rule of requiring that there be 2 or more events within the first 15 events that have means >80 was sufficient to automatically separate the files this way for visualization.

##### Looking at the number of “0 moves” (stays) vs. length and/or quality

The fragstats.txt file produced above was brought into R^1^ for the various plots comparing the number of stays (“0 moves”) with other features such as read length, number of events, number of called events, and mean quality scores (Q).

##### Identifying G4 motif positions in template and complement reads

The poreminion subcommand “g4” was used in the following way:

$ poreminion g4 –minG 3 –maxN 7 –numtracts –noreverse -f5 all/ > g4s.bed

This subcommand uses the quadparser^3,4^ regular expression, G_3+_-N_1-7_G_3+_N_1-7_G_3+_N_1-7_G_3+_ (regular expression in Python: ‘([gG]{3,}\w{1,7}){3,}[gG]{3,}’) to search the sequences inside fast5 files for G4 motifs (G_3+_ is specified by “—minG 3” and N_1-7_ is specified by “—maxN 7”). The “—noreverse” option specifies to only search the sequence given (not its complement), or in other words it specifies to NOT also search for the “C4” motif: C_3+_-N_1-7_C_3+_N_1-7_C_3+_N_1-7_C_3+_. The “—numtracts” option reports the number of poly-G tracts inside a given G4 motif as the Python regular expression (above) searches for 4 or more adjacent poly-G tracts separated by 1-7 nucleotides. The more poly-G tracts, the more possible ways a G4 structure could form. For example, observing five poly-G tracts does not simply indicate two overlapping G4 motifs that can form only two G4 structures, but rather “5 choose 4” (5) ways to choose four of those five poly-G tracts multiplied by the number of ways 4 poly-G tracts can fold together into a G4 structure (e.g. there are parallel and anti-parallel arrangements) as well as the multiplicative possibilities when varying the number and position of consecutive Gs (≥3) used in the chosen poly-G tracts. Keeping track of the number of poly-G tracts inside each G4 motif allowed us to separate the G4 motifs into two groups: those with 4 poly-G tracts and those with >4 for the subsequent statistical analysis testing which group has more “0 moves” associated with it on average.

##### Identifying positions of stays (“0 moves”) in template and complement reads

The poreminion subcommand “staypos” was used as follows:

$ poreminion staypos all/ > stays.bed

This subcommand goes through the base-called events of template and complement strands while keeping track of the index of each event relative to the output sequence (by accounting for base-caller “moves” of 0-5) and reporting the positions in the sequence that correspond to “0 moves” in the base-called events. Specifically, it reports the coordinates of the 5mer corresponding to the “0 move” in BED format (for example, if a 5mer starts at position 0, its end position is 5).

##### Comparing G4 and Stay positions in template and complement reads

For plotting, windowBed from BEDtools^5^ was used to obtain all stay positions within 500 nucleotides of a G4 motif (done independently for each of the three runs):

$ windowBed -a g4s.bed -b stays.bed -w 500 > g4s.stays.500.windowbed

The resulting file contains lines with pairs of entries for the G4 motif position and stay (“0 move”) position for each pair that is within 500 nucleotides of each other. This file was brought into R^1^ where distances between G4 centers and ‘stay’ centers (i.e. the middle nucleotide of a 5mer) from G4-stay pairs were calculated as the distance between their centers. Centers were found by subtracting 1 from the end position of each BED entry (to account for BED format), then taking the mean of the start and resulting end positions. Histogram information was obtained by, for example, hist(distances, breaks=seq(from=-650.5, to=650.5, by=1), which results in the histogram midpoints being integers from -650 to 650. Histogram counts were lightly loess smoothed (loess(hist.counts ∼ hist.mids, span=0.05).

Four null distributions were considered – for each G4 motif, a site of the same length was selected uniformly at random from: (null 1) any template or complement read with no G4 motifs, (null 2) any template or complement read, (null 3) anywhere within the same read the G4 motif was on, (null 4) anywhere within the same read the G4 motif was that did not overlap the G4 motif coordinates. These coordinates were selected with BEDtools^5^:

$ shuffleBed -noOverlapping -excl readswithg4.bed -i g4s.bed -g template-and-complement-reads > g4s.shuffled.nonG4reads.bed

$shuffleBed -noOverlapping -i g4s.bed -g template-and-complement-reads > g4s.shuffled.allreads.bed

$shuffleBed -noOverlapping -chrom -i g4s.bed -g template-and-complement-reads > g4s.shuffled.sameread.bed

$shuffleBed -allowBeyondChromEnd -noOverlapping -chrom -excl g4s.bed -i g4s.bed -g template-and-complement-reads > g4s.shuffled.sameread.notoverG4.bed

windowBed from BEDtools was used as above to collect pairs of randomly selected locations and “0 moves” that were within 500 nucleotides of each other. Histogram counts and smoothing was same as above. These null distributions were plotted on same plots as above.

For statistical analyses, we compared the number of “0 moves” within 50 nucleotides of the G4 motifs with their matched null positions from the four different null distributions. These counts were obtained using windowBed from BEDtools^5^. For example:

windowBed -c -a g4s.bed -b stays.bed -l 50 -r 50 > g4s.stays.50.counts.txt

windowBed -c -a nulls.bed -b stays.bed -l 50 -r 50 > nulls.stays.50.counts.txt

For each of the four nulls described above, the pairs of counts from G4 motifs and the null were used as input to the “sign test” as well as the Wilcoxon signed rank test in R^1^. Note that there seems to be more “0 moves” on reads with G4 motifs in general. The fourth null (null 4: selecting a random position on the same read as the G4 motif that does not overlap the G4 motif) serves as the best matched pairs, controlling for any read-specific effects and still all p-values were significant (Supplementary Table 6A). To test the hypothesis that G4 motifs on the complement strand associated with more “0 moves” than G4 motifs on the template strand, the counts for each of these group was used as input to the Wilcoxon rank sum test in R^1^ (Supplementary Table 6B). To test the hypothesis that G4 motifs with >4 poly-G tracts were associated with higher “0 move” counts than G4 motifs with only 4 poly-G tracts, the counts for each of these groups was used as input to the Wilcoxon rank sum test in R^1^ (Supplementary Table 6C).

### Supplementary Figures

**Supplementary Figure 1:**
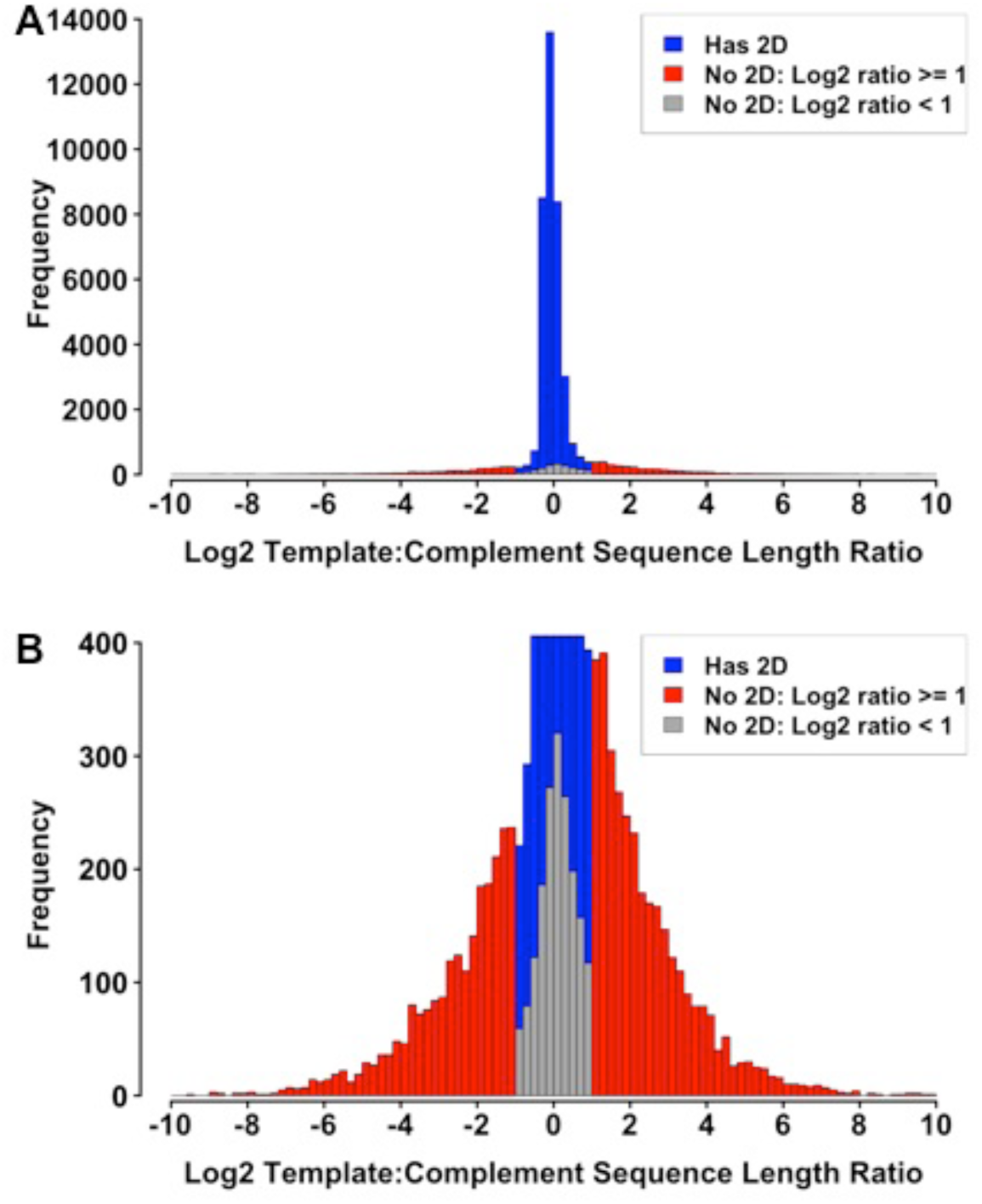
Distribution of Log2(template:complement) for base-called fast5 files that contain both template and complement reads. **(A)** shows a zoomed-out view of a histogram of Log2(ratio of template read length to complement read length) for all base-called fast5 files that have both template and complement reads. The histogram shows the number (frequency) of fast5 files as a function of Log2(ratio). **(B)** shows a zoomed-in view of the bottom of the same histogram as in A. Most fast5 files that have both template and complement reads also have 2D reads (blue). For those that do not, most have template-to-complement read length ratios that are either too big (>2, or >1 in log2) or too small (<0.5, or <-1 in log2) for initiating 2D base-calling (red). However, there are also base-called fast5 files with both template and complement reads where the ratio is within range for 2D base-calling and where 2D base-calling fails (grey).

**Supplementary Figure 2:**
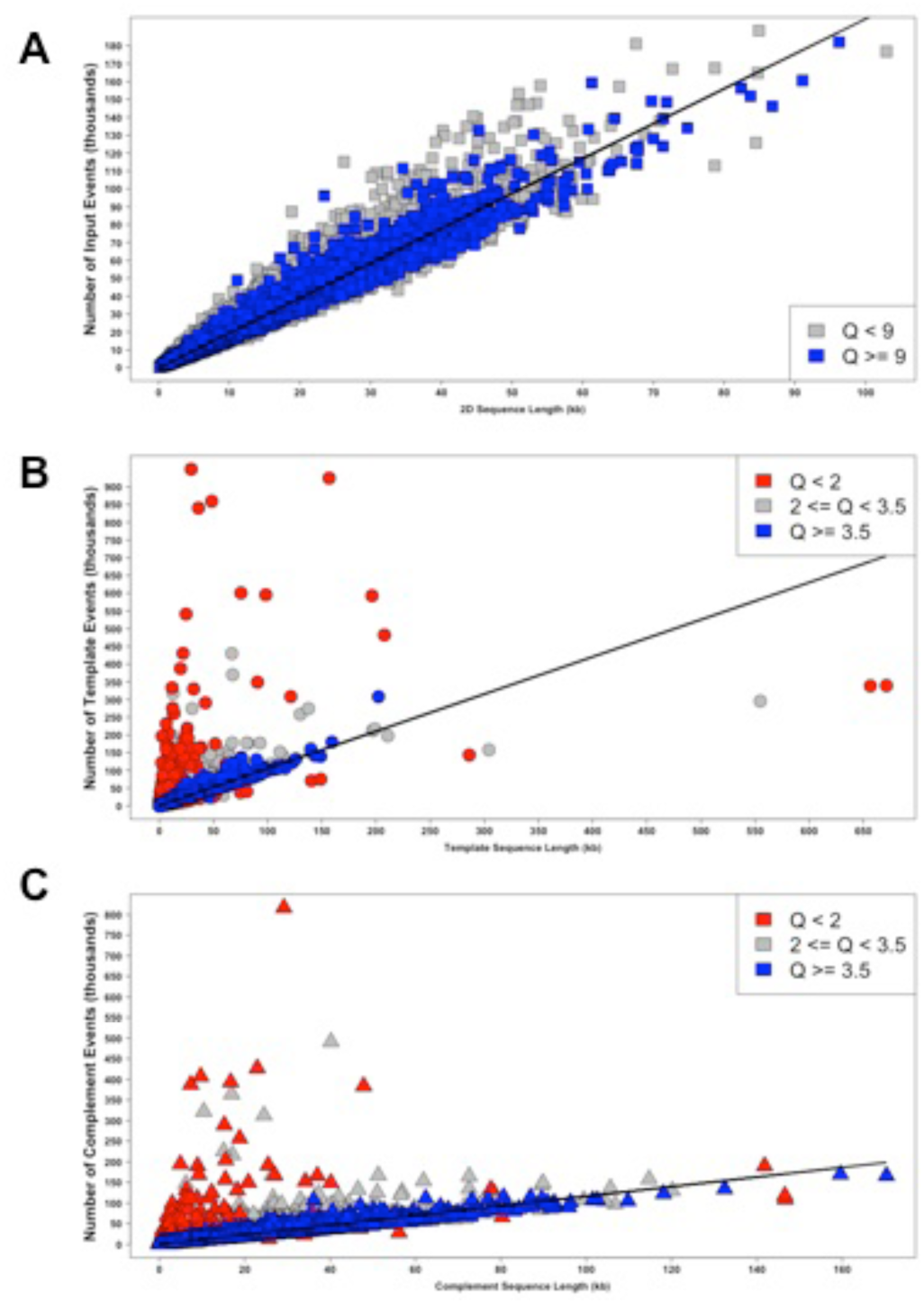
Number of pre-base-calling events vs. post-base-calling sequence length. **(A)** Number of total input events vs. 2D read length for reads with 2D base-calls. The slope of line is approximately 2 (1.95) representing ∼2 events per nucleotide on average for fast5 files that contain a 2D read. Blue represents the high quality (Q≥9) 2D reads. Number of template events (not to be confused with number of called template events where some are pared out) versus template sequence length. The slope (1.051) of ∼1 indicates ∼1 event per nucleotide on average. Blue represents higher quality 1D reads (Q≥3.5) while red represents lower quality 1D reads (Q<2). **(C)** Number of complement events versus complement sequence lengths shows a slope of ∼1 (1.166) representing ∼1 event per nucleotide on average. Blue and red are as in B. Prior to base-calling, one can estimate molecule size using 1-2 events per nucleotide (depending on whether the fast5 file contains only template events, if there are also complement events, and if so, how many) noting that a minority of exceptions have higher event:nucleotide ratios. This means that 1-2 events per nucleotide is often an upper limit estimate for molecule size. Importantly, template and complement sequences with mean quality scores ≥3.5 (blue in B and C) stay tightly packed around the ∼1 event:nt line whereas those with Q<3.5 (grey and red in B and C) often fall off the line (especially when Q<2; red), demonstrating that using Q≥3.5 as a cut-off for 1D reads has a practical interpretation that may be useful.

**Supplementary Figure 3:**
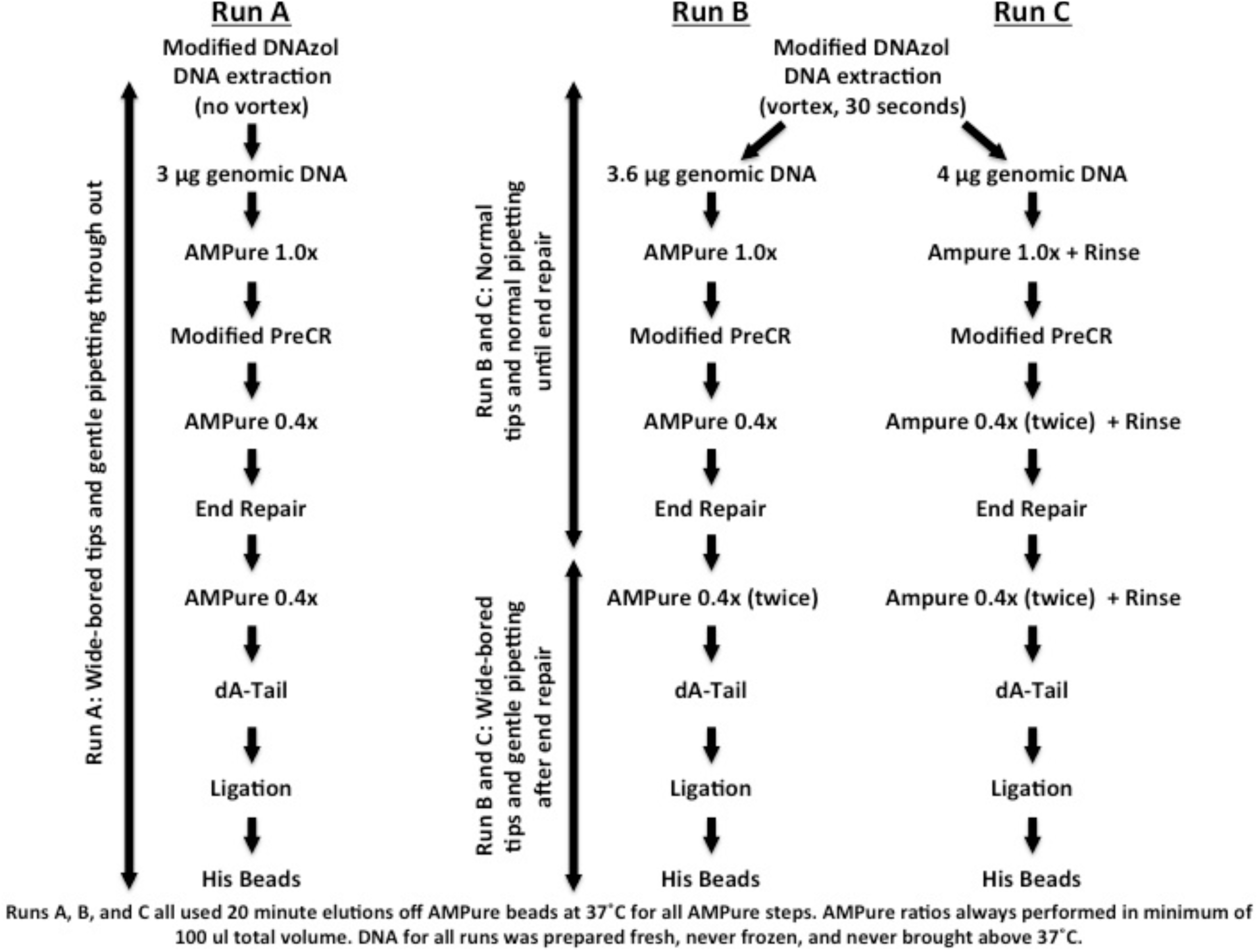
See Supplementary Methods for more information. Briefly, for **Run A**, we gently let freshly obtained precipitated DNA re-suspend in TE (pH 8.0), skipped the Covaris shearing step in the standard protocol, used wide-bored tips with gentle pipetting to minimize DNA breakage, and started with 3X the recommended starting material (3 μg instead of 1μg) to compensate for differences in molarity. PreCR was done in double the volume with double the reagents for double the time since we started with more DNA than recommended. Both AMPure steps after PreCR were done at 0.4x. For **Run B**, we vortexed the DNA (full speed, 30 seconds) after DNA extraction and used normal pipette tips until end-repair, after which wide-bored tips and gentler pipetting were employed. Moreover, to account for the possible increase of molecules <1kb due to vortexing, we started with more material (3.6x) and performed two sequential 0.4x AMPure bead clean-ups after end-repair. For **Run C**, we used 4x the recommended input amount of DNA, did two sequential 0.4x AMPure clean-ups before and two after end-repair, and did a new rinse step (see Supplementary Methods and Supplementary Fig 4B) at the end of all AMPure steps before the final elution of DNA off the beads. In the rinse, smaller DNA preferentially falls off the beads. For all runs, in all AMPure elutions, the beads were incubated at 37°C for ≥20 minutes to facilitate and wait for long DNA to come off the beads. Also, DNA was never subject to temperature extremes below 4°C or above 37°C and was re-suspended in 1X TE, pH 8 when isolated. Moreover, in AMPure bead steps, the ethanol washes were performed with buffered 80% ethanol (10 mM Tris-Cl, pH 8).

**Supplementary Figure 4:**
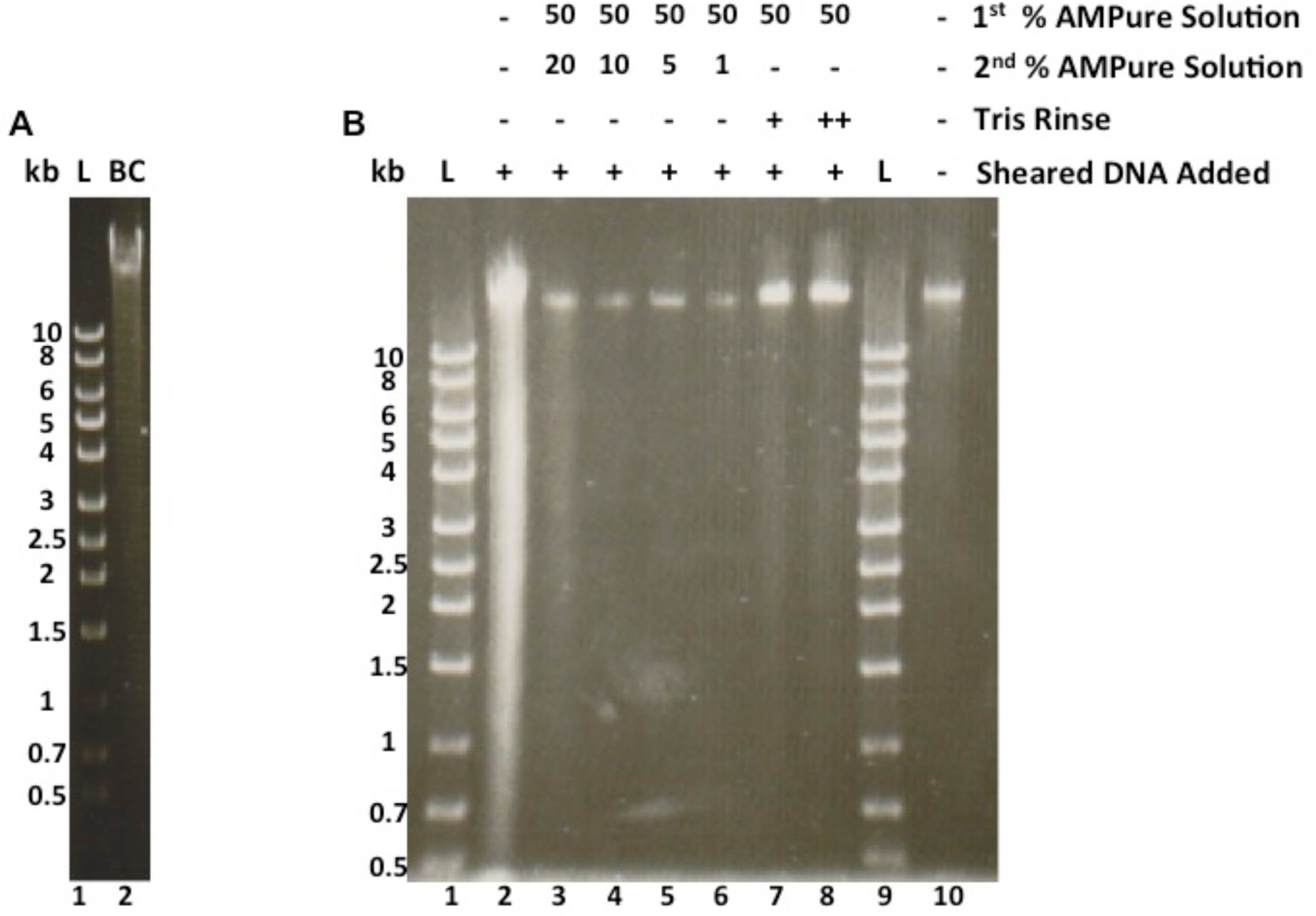
**(A)** “L” is the DNA ladder and “kb” shows the sizes of the bands in the ladder. “BC” is the genomic DNA used for both Run B and Run C. Vortexing was used to help re-suspend the DNA after extraction. This gel shows that a substantial proportion remains above 10 kb nonetheless. **(B)** “L” and “kb” same as in A. Genomic DNA was gently extracted and re-suspended. Eight 15 μl aliquots were set aside. The rest was lightly sonicated/sheared using the BioRuptor (Diagenode) to a very broad range with a lower limit near 0.5kb and an upper limit at the size of unsheared DNA (unsheared DNA shown in lane 10). Equal volumes of sheared DNA were added to 7 of the 8 aliquots (lanes 2-8, not lane 10). We then tried several strategies to deplete the shorter DNA (e.g. <10kb), all using AMPure beads (Lanes 3-8). Lane 2 shows the sheared+unsheared DNA without depletion of smaller DNA. Each of the aliquots with sheared DNA added (Lanes 2-8) was brought to 100 μl volume with Ultra Pure Water (UPW, Life Technologies UltraPure DNase/RNase-free, distilled water). 100 μl AMPure beads were then added to each to create a 50% AMPure mixture (or 1.0x ratio). The DNA was incubated in the 50% AMPure solution for 5 minutes before pelleting the beads on a magnet for 5 minutes and removing the supernatant. The beads were then washed twice with 80% ethanol while on the magnet. After the second 80% ethanol wash, the tubes with the beads were lightly spun and placed on the magnetic rack for 1 minute before removing the remaining 80% ethanol collected at the bottom from the light spin. The previous steps were performed to remove any buffers or salts associated with the DNA to have finer control over the AMPure ratio (for lanes 3-6) in subsequent steps. The beads from lane 2 were air-dried and eluted at this point. For lanes 3-8, instead of eluting, the beads were re-suspended in a second AMPure solution or “rinsed” once (lane 7) or twice (lane 8) before eluting. For lanes 3-6, AMPure solutions were premade by combining 40 μl AMPure with 160 μl UPW (20%, lane 3), 20 μl AMPure with 180 μl UPW (10%, lane 4), 10 μl AMPure with 190 μl UPW (5%, lane 5), and 2 μl AMPure with 198 μl UPW (1%, lane 6). In all cases (lanes 3-6), the second AMPure solution was added to the beads from the first AMPure step (after the 80% ethanol washes described above), and the beads were gently re-suspended, incubated for 10 minutes, and put on a magnetic rack for 5 minutes before the supernatant was removed and two 80% ethanol washes were performed while the beads remained on the rack. After the second 80% ethanol wash, the tubes with the beads were lightly spun and placed on the magnetic rack for 1 minute before removing the remaining 80% ethanol collected at the bottom from the light spin. The tubes were then allowed to air dry for 2-3 minutes. For lanes 7-8, after the first set of 80% ethanol washes, instead of proceeding to a second AMPure step, the tubes with the beads were lightly spun and placed on the magnetic rack for 1 minute before removing the remaining 80% ethanol collected at the bottom from the light spin. The tubes were then allowed to air dry on the magnetic rack for 2 minutes before adding 200 μl UPW very gently to the tube-wall opposite the beads while they remained on the magnetic rack. This is the “rinse”. The beads were allowed to incubate in the rinse for 1 minute before very gently removing it. Lane 8 was subject to a second identical rinse, which seems to offer minimal additional benefits beyond the first rinse (compare lanes 7 and 8). For all elutions (lanes 2-8), DNA was eluted off the AMPure beads into 15 μl UPW at 37°C for 20 minutes to facilitate elution of long DNA. Loading buffer was added to each sample (lanes 2-8,10). The DNA samples were then loaded onto the gel, electrophoresed, and stained with ethidium bromide for visualization. Although all conditions eliminated most of the smaller DNA (e.g. <10 kb), it was possible to tell where the tails of the smears in each lane ended using longer exposure times (data not shown). Lane 3 (20% AMPure) ended around 1 kb, Lane 4 (10% AMPure) ended above 2 kb, Lane 5 (5% AMPure) ended above 3 kb, Lane 6 (1% AMPure) ended above 5 kb, Lane 7 (1 rinse) ended above 1 kb, and Lane 8 (2 rinses) ended above 1.5 kb. While some of the lower percentage AMPure solutions performed better than the “rinse” as judged by where the tail ended, there were also comparable losses of the large DNA (compare lanes 3-6 to lane 2). In contrast, the rinses largely removed the small DNA while retaining the majority of the large DNA (compare lanes 7-8 with lane 2). These results were used to inform us how to modify the AMPure steps for Run C. To deplete DNA <10kb in Run C, we chose to (1) keep the strategy of sequential AMPure washes as done here (Lanes 3-6), but with the ONT recommended 0.4x AMPure ratio (∼28-29% AMPure solution) in the first <u>and</u> second sequential steps instead of the 50% (1x) AMPure solution used in the first step and lower percent (1-20%) solutions used in the second step here and (2) add a rinse step after the second sequential AMPure step. This was successful as determined by comparing the read lengths from Runs B and C, which came from the same source of DNA (Fig. 1D, Supplementary Fig. 5).

**Supplementary Figure 5:**
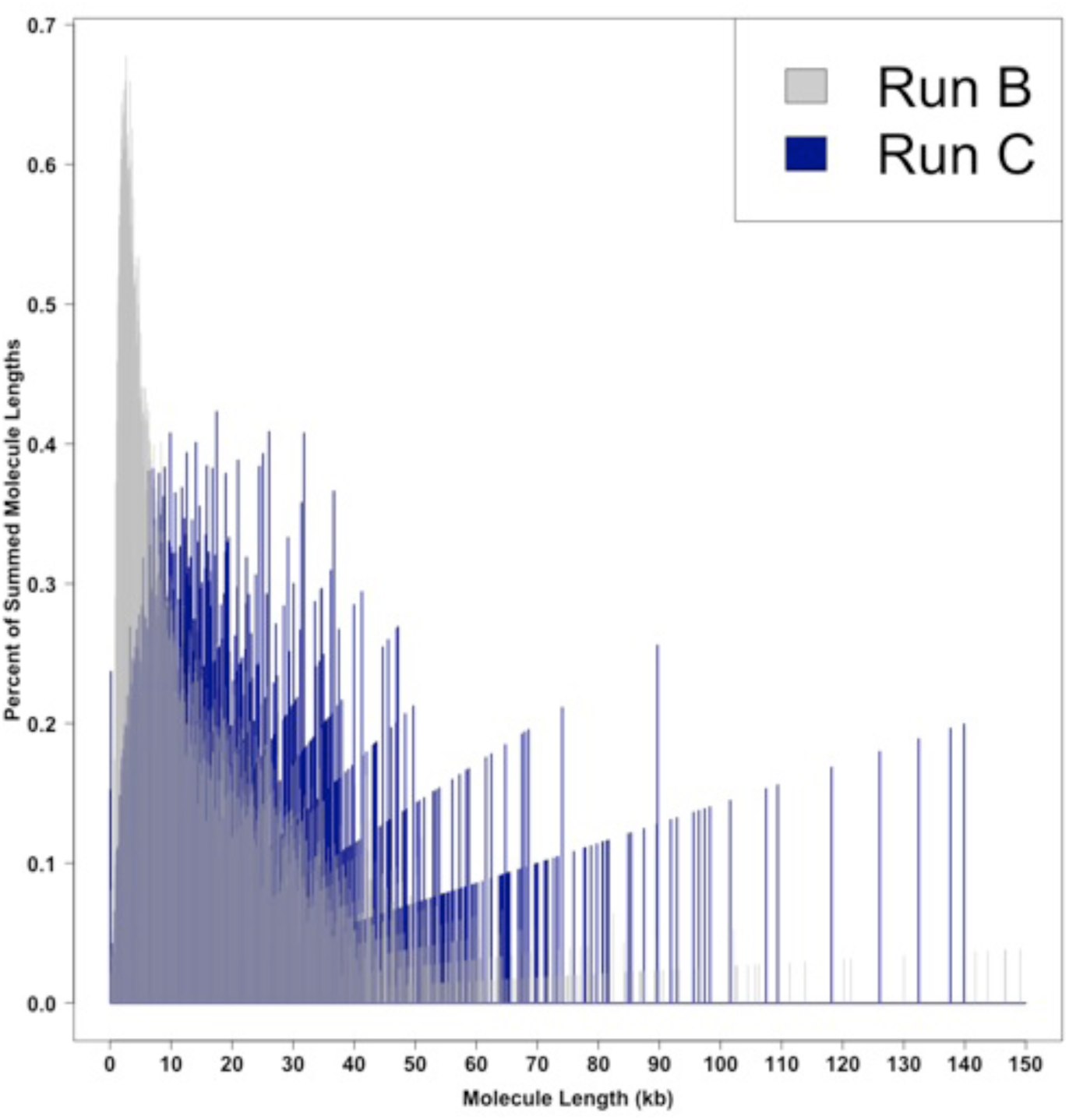
The proportion of total summed molecule length as a function of molecule length for Run B (transparent grey) and Run C (dark blue). Note that the darker shade of grey is a result of the dark blue behind the transparent grey. Run B and Run C used the same source of DNA, but differed in library preparation (see Supplementary Fig. 3). Run C used a new rinse step during all AMPure clean-ups as well as took additional advantage of sequential AMPure rounds in each clean-up step.

**Supplementary Figure 6:**
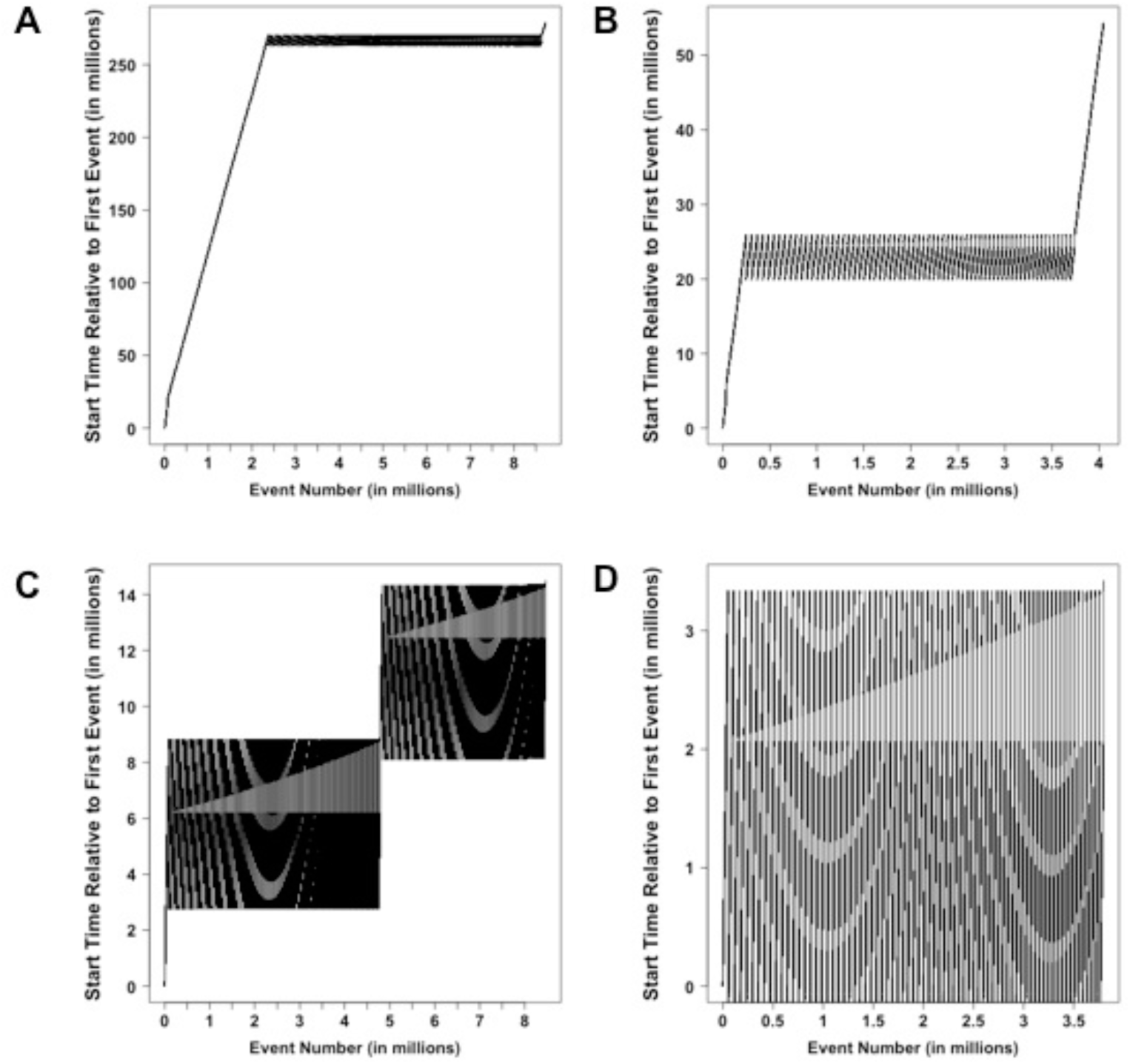
Examples of multi-million event files that contained “Time Errors” (repeated blocks of events). **(A)** Example from Run A. **(B)** Another example from Run A. **(C)** Example from Run B. **(D)** Example from Run C. The “event start time” as a function of “event number” should be a monotonically increasing function. However, in files with repeated blocks of events, the function is not monotonically increasing and instead exhibits periodic patterns.

**Supplementary Figure 7:**
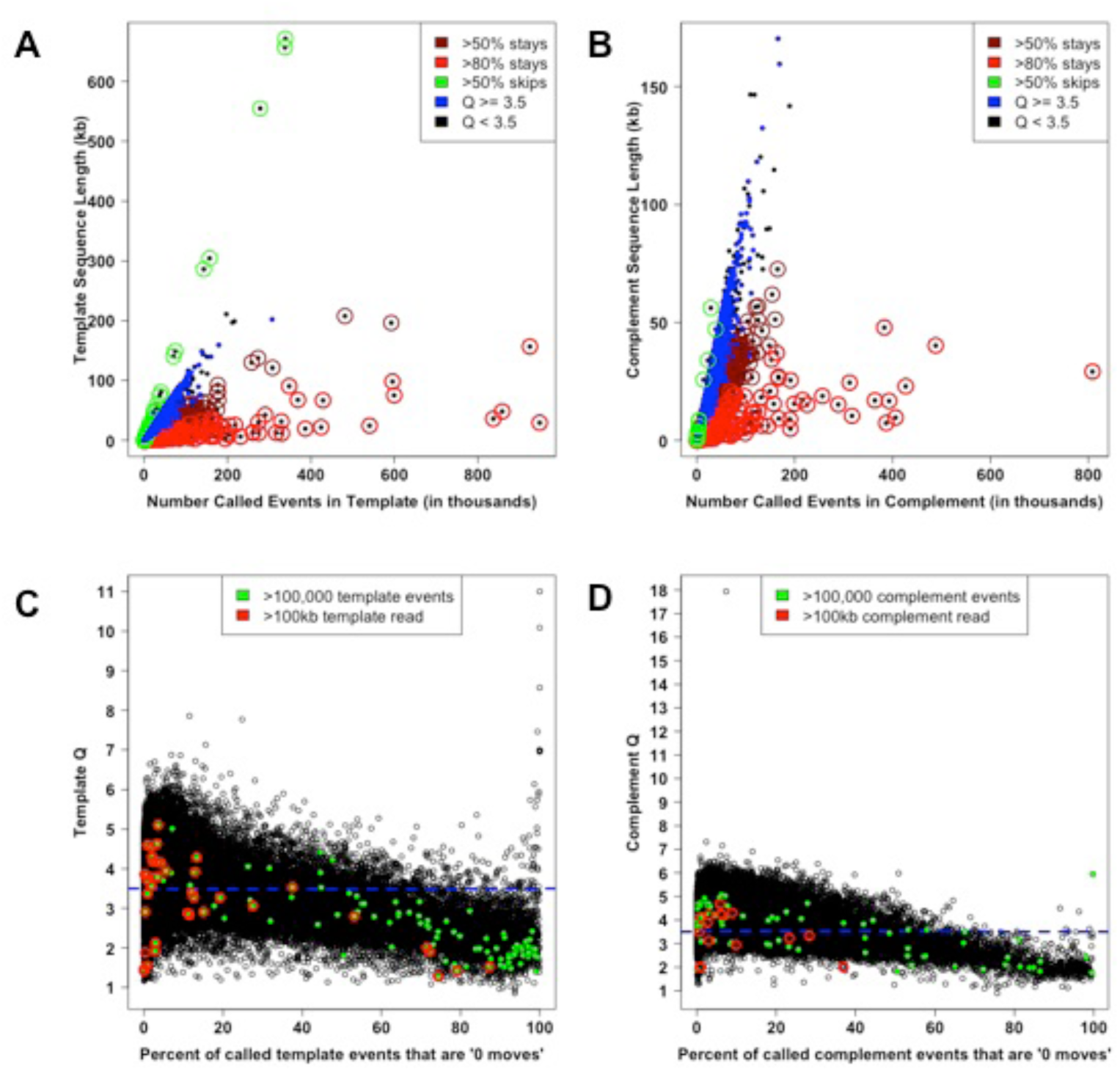
Number of base-called events, sequence lengths, percent of base-called events assigned “0 moves”, and mean quality scores. **(A)** Number of base-called template events vs. template sequence length with information on mean quality scores and percent of base-called template events assigned a “0 move” (“stay”). **(B)** Number of base-called complement events vs. complement sequence length with information on mean quality scores and percent of base-called complement events assigned a “0 move” (“stay”). **(C)** Percent of called template events assigned a “0 move” (stay) vs. mean quality score (Q). Slope of best fit line (lm(y ∼ x) in R) is -0.01713 meaning for every 1 percent (or 10 percent) increase in “0 moves” there is an average decrement of 0.01713 (or 0.1713) in Q. **(D)** Percent of called complement events assigned a “0 move” (stay) vs. mean quality score (Q). Slope of best fit line (lm(y ∼ x) in R) is -0.02363 meaning for every 1 percent (or 10 percent) increase in “0 moves” there is an average decrement of 0.02363 (or 0.2363) in Q. In (A) and (C), higher quality 1D reads (Q≥3.5) are blue and those less than that are black. Maroon circles are around points with ≥50% of base-called events assigned a “0 move”, red circles are around points with ≥80% of base-called events assigned a “0 move”, and green circles are around points where ≥50% of the base-called events were “skips” (move=2). In (C) and (D), the blue dashed line is at Q=3.5, large green points represent points with >100,000 called template (C) or complement (D) events and maroon circles encompass points that had ≥100 kb template (C) or complement (D) sequences.

**Supplementary Figure 8:**
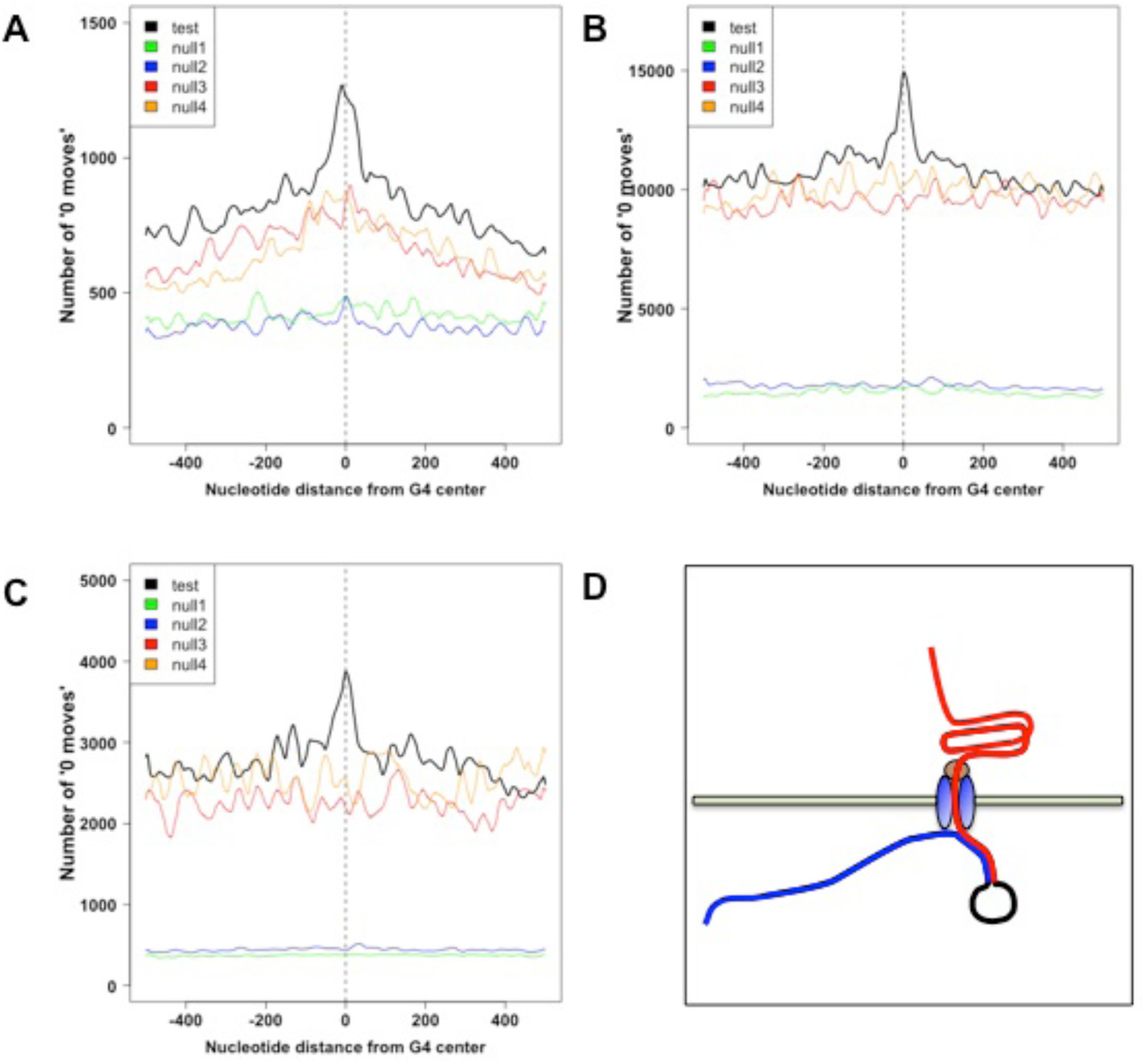
Aggregate analyses of the distribution of “0 moves” around G4 motif centers. The number of “0 moves” as a function of proximity to G4 motif centers (test condition) or the centers of randomly selected positions (null 1-4 conditions) for **(A)** Run A, **(B)** Run B, and **(C)** Run C. **(D)** Depicts the orientation (5’ – 3’) of a DNA molecule going through a pore from the top chamber to the bottom chamber and how a G4 might cause a DNA molecule to stall resulting in an accumulation of measurements of a 5mer or set of 5mers slightly upstream of the G4 (the adjacent upstream sequence is pulled through the pore before the downstream G4 blocks further translocation). The template strand is blue, the hairpin is black, and the complement strand is red.

### Supplementary Tables

**Supplementary Table 1A:** Statistics on Mean Quality Scores (Q) for 2D and 1D reads from Run A.

| Run A | 2D | 1D | Template | Complement |
| --- | --- | --- | --- | --- |
| **Median Q** | 9.23666606763 | 3.54781658091 | 3.41144557734 | 4.0100659241 |
| **Mean Q** | 9.13260594983 | 3.553805971 | 3.43220450158 | 3.93551548089 |
| **Standard Deviation of Q** | 1.08997018784 | 0.916461481548 | 0.871848558243 | 0.947880924972 |
| **Minimum Q** | 5.79424190524 | 0.965373861239 | 0.965373861239 | 1.12938538308 |
| **Maximum Q** | 12.2263387187 | 10.0869987652 | 10.0869987652 | 6.43558322934 |

The group of 1D reads contains all template and complement reads.

**Supplementary Table 1B:**
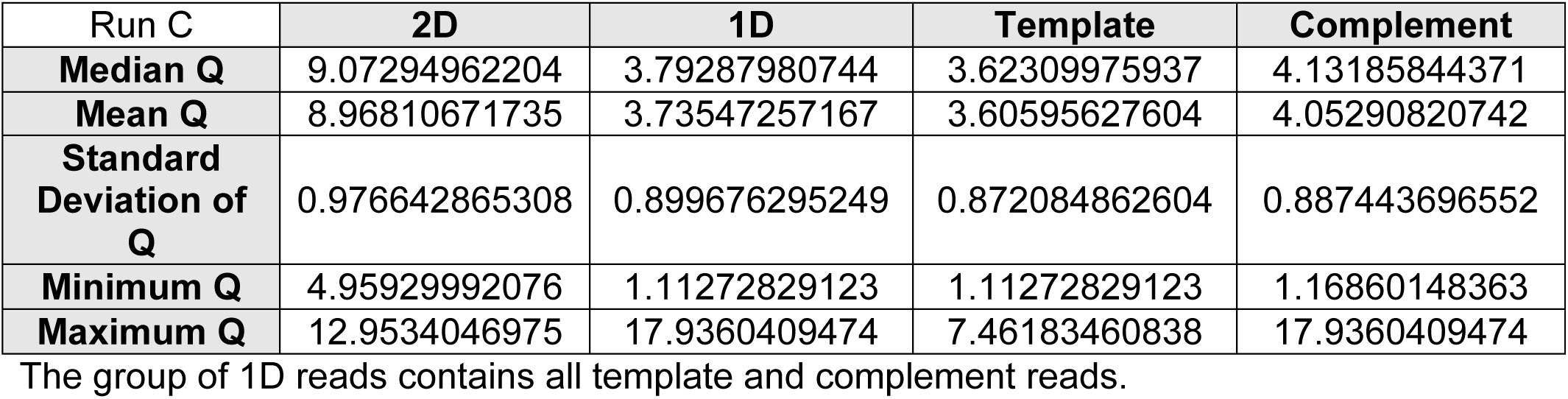
Statistics on Mean Quality Scores (Q) for 2D and 1D reads from Run B.

**Supplementary Table 1C:**
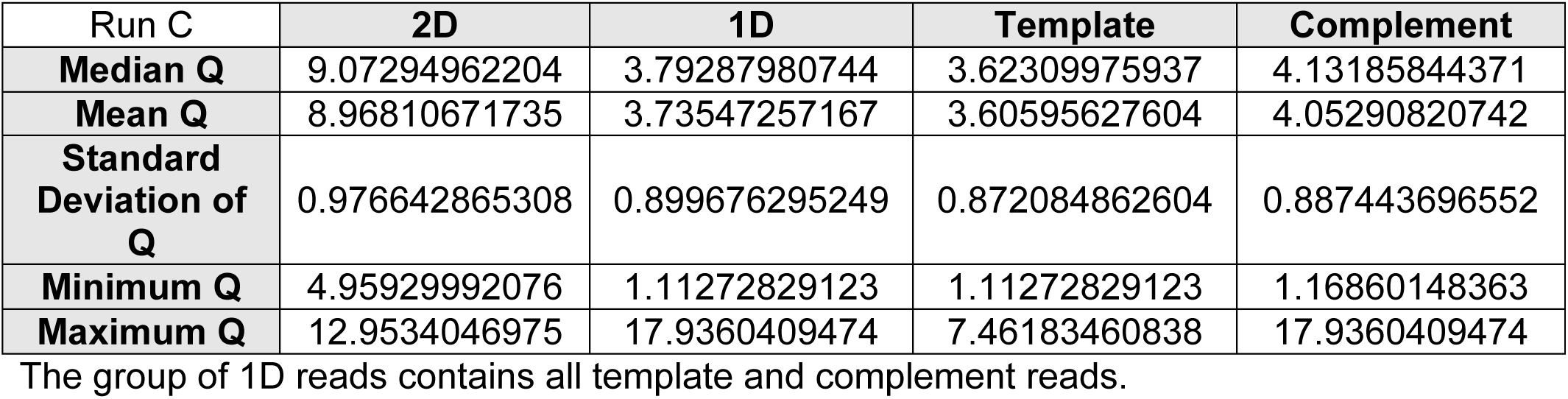
Statistics on Mean Quality Scores (Q) for 2D and 1D reads from Run C.

**Supplementary Table 2:** Read types in base-called fast5 files.

|  | **Run**<br>**A** | **Run**<br>**B** | **Run**<br>**C** |
| --- | --- | --- | --- |
| n_molecules (# basecalled files) | 9935 | 64282 | 8941 |
| n_template_only | 6770 | 26923 | 5293 |
| percent_template_only | 68.14 | 41.88 | 59.20 |
| n_has_complement | 3165 | 37359 | 3648 |
| percent_has_complement | 31.86 | 58.12 | 40.80 |
| n_has_2D | 2172 | 31999 | 2523 |
| percent_has_2D | 21.86 | 49.78 | 28.22 |
| n_does_NOT_have_2D | 7763 | 32283 | 6418 |
| percent_does_NOT_have_2D | 78.14 | 50.22 | 71.78 |
| percent_of_molecules_that_do_NOT_have_2D<br>because_template_only | 87.21 | 83.40 | 82.47 |
| percent_of_molecules_that do_NOT_have_2D_that<br>DO_have_complement | 12.79 | 16.60 | 17.53 |
| n_with_complement_that_DO_have_2D | 2172 | 31999 | 2523 |
| n_with_complement_that do NOT_have_2D | 993 | 5360 | 1125 |
| percent_of_molecules_with_complement that_DO_have_2D | 68.63 | 85.65 | 69.16 |
| percent_of_molecules_with_complement that do NOT_have_2D | 31.37 | 14.35 | 30.84 |

**Supplementary Table 3:** Molecule and Read Length Statistics. All lengths are nucleotide lengths in Supplementary Tables 3A-E (e.g. 10,000 = 10,000 nucleotides).

**Supplementary Table 3A:** Molecule Statistics.

|  | Run A | Run B | Run C |
| --- | --- | --- | --- |
| sum_molecule_lengths | 49,778,211 | 386,880,692 | 70,086,870 |
| Molecule N25 | 46,973 | 28,006 | 34,939 |
| Molecule N50 | 25,238 | 13,553 | 20,824 |
| Molecule N75 | 11,969 | 5,279 | 10,749 |
| mean_molecule_size | 5,010.4 | 6,018.5 | 7,838.8 |
| median_molecule_size | 152 | 2,817 | 3,116 |
| max_molecule_size | 304,309 | 671,219 | 139,864 |
| min_molecule_size | 5 | 5 | 5 |
| n_molecules_gt_10kb | 1,433 | 10,145 | 2,318 |
| percent_molecules_gt_10kb | 14.42 | 15.78 | 25.93 |
| sum_molecules_gt_10kb | 39,415,998 | 226,507,295 | 54,080,719 |
| percent_summed_molecules_from_molecules_gt_10kb | 79.18 | 58.55 | 77.16 |
| n_molecules_gt_50kb | 139 | 396 | 119 |
| percent_molecules_gt_50kb | 1.40 | 0.62 | 1.33 |
| sum_molecules_gt_50kb | 10,999,924 | 27,504,214 | 8,168,033 |
| percent_summed_molecules_from_molecules_gt_50kb | 22.10 | 7.11 | 11.65 |
| n_molecules_gt_100kb | 21 | 24 | 8 |
| percent_molecules_gt_100kb | 0.21 | 0.04 | 0.09 |
| sum_molecules_gt_100kb | 3,140,720 | 4,779,702 | 972,486 |
| percent_summed_molecules_from_molecules_gt_100kb | 6.31 | 1.24 | 1.39 |

Molecule size is determined as (1) the length of the 2D read if one is available, (2) the length of the template if only the template is available, or (3) the longer of the template and complement reads when both are available in the absence of a 2D read (See supplementary methods). Thus, the statistics are performed on a set of unique/non-redundant molecule length estimates (a single read for each molecule as opposed to 1-3 reads per molecule).

**Supplementary Table 3B:**
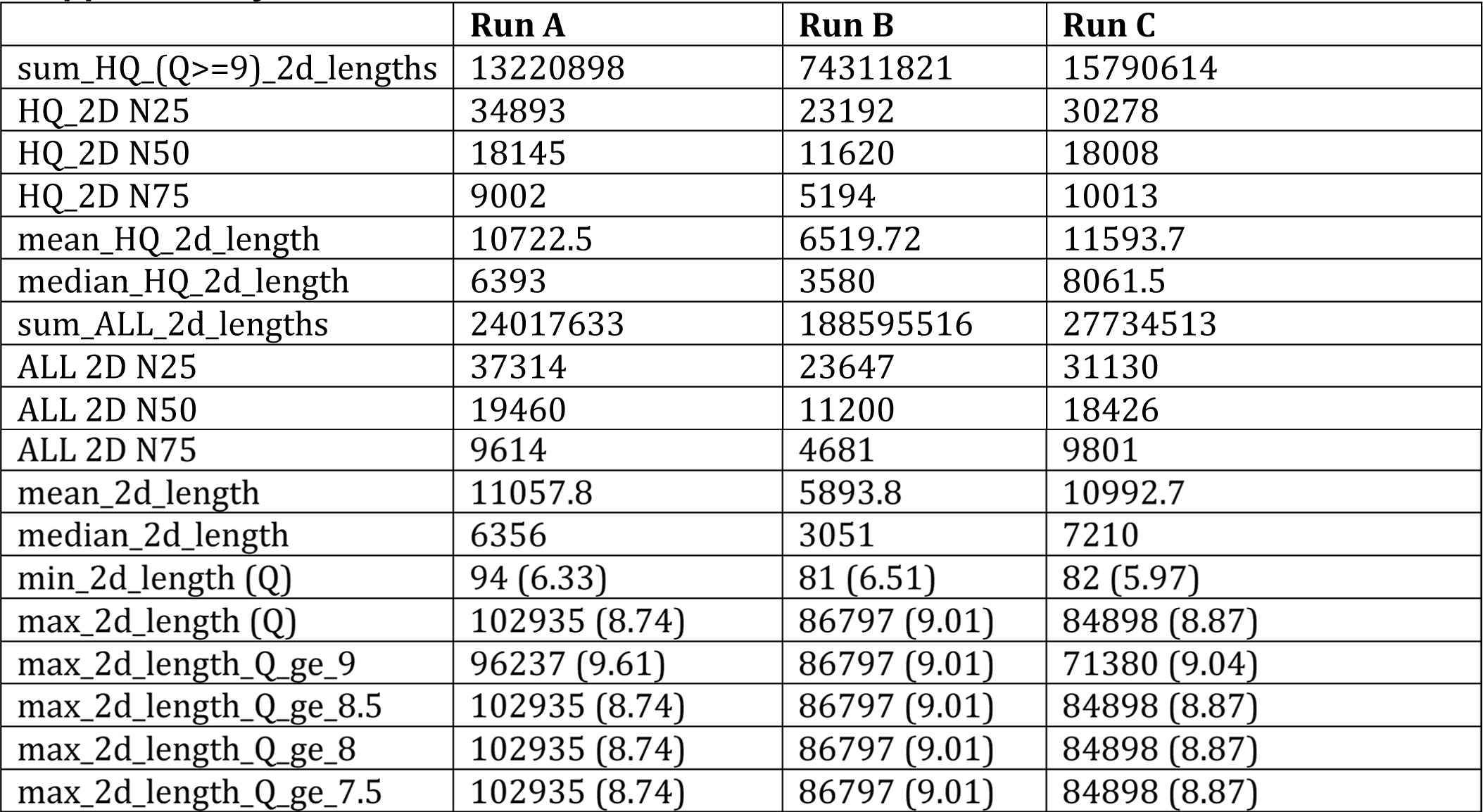
2D Read Statistics.

|  | Run A | Run B | Run C |
| --- | --- | --- | --- |
| sum_HQ_(Q>=9)_2d_lengths | 13220898 | 74311821 | 15790614 |
| HQ_2D N25 | 34893 | 23192 | 30278 |
| HQ_2D N50 | 18145 | 11620 | 18008 |
| HQ_2D N75 | 9002 | 5194 | 10013 |
| mean_HQ_2d_length | 10722.5 | 6519.72 | 11593.7 |
| median_HQ_2d_length | 6393 | 3580 | 8061.5 |
| sum_ALL_2d_lengths | 24017633 | 188595516 | 27734513 |
| ALL 2D N25 | 37314 | 23647 | 31130 |
| ALL 2D N50 | 19460 | 11200 | 18426 |
| ALL 2D N75 | 9614 | 4681 | 9801 |
| mean_2d_length | 11057.8 | 5893.8 | 10992.7 |
| median_2d_length | 6356 | 3051 | 7210 |
| min_2d_length (Q) | 94 (6.33) | 81 (6.51) | 82 (5.97) |
| max_2d_length (Q) | 102935 (8.74) | 86797 (9.01) | 84898 (8.87) |
| max_2d_length_Q_ge_9 | 96237 (9.61) | 86797 (9.01) | 71380 (9.04) |
| max_2d_length_Q_ge_8.5 | 102935 (8.74) | 86797 (9.01) | 84898 (8.87) |
| max_2d_length_Q_ge_8 | 102935 (8.74) | 86797 (9.01) | 84898 (8.87) |
| max_2d_length_Q_ge_7.5 | 102935 (8.74) | 86797 (9.01) | 84898 (8.87) |

**Supplementary Table 3C:** 1D Read Statistics.

|  | Run A | Run B | Run C |
| --- | --- | --- | --- |
| sum_1d_lengths | 75870350 | 565275221 | 102225612 |
| 1D N25 | 41287 | 25353 | 33308 |
| 1D N50 | 21927 | 12037 | 19812 |
| 1D N75 | 10432 | 4776 | 10349 |
| mean_1d_length | 5791.6 | 5561.5 | 8120.2 |
| median_1d_length | 231 | 2661 | 3783 |
| sum_template_lengths | 43984348 | 349320608 | 66858648 |
| Template N25 | 43176 | 25699 | 33509 |
| Template N50 | 22324 | 12198 | 20001 |
| Template N75 | 10344 | 4752 | 10421 |
| mean_template_length | 4427.2 | 5434.2 | 7477.8 |
| median_template_length | 151 | 2565.5 | 2892 |
| min_template_length (Q) | 5 (10.09) | 5 (6.95) | 5 (6.97) |
| max_template_length | 304309 (2.12) | 671219 (1.55) | 139864 (4.28) |
| max_template_Q_ge_4_length | 122689 (4.63) | 143763 (4.17) | 139864 (4.28) |
| max_template_Q_ge_3.5_length | 202293 (3.53) | 143763 (4.17) | 139864 (4.28) |
| max_template_Q_ge_3_length | 210931 (3.36) | 143763 (4.17) | 139864 (4.28) |
| max_template_Q_ge_2.5_length | 210931 (3.36) | 554720 (2.91) | 139864 (4.28) |
| sum_complement_lengths | 31886002 | 215954613 | 35366964 |
| Complement N25 | 39391 | 24784 | 33054 |
| Complement N50 | 21394 | 11821 | 19287 |
| Complement N75 | 10506 | 4825 | 10163 |
| mean_complement_length | 10074.6 | 5780.5 | 9694.9 |
| median_complement_length | 5049 | 2836 | 5700 |
| min_complement_length | 17 (1.13) | 14 (3.63) | 10 (4.36) |
| max_complement_length | 170334 (3.87) | 146631 (2.00) | 132439 (4.20) |
| max_complement_Q_ge_4_length | 159563 (4.30) | 102671 (4.20) | 132439 (4.20) |
| max_complement_Q_ge_3.5_length | 170334 (3.87) | 102671 (4.20) | 132439 (4.20) |
| max_complement_Q_ge_3_length | 170334 (3.87) | 105683 (3.23) | 132439 (4.20) |
| max_complement_Q_ge_2.5_length | 170334 (3.87) | 120149 (2.94) | 132439 (4.20) |

Note: Statistics on “1D reads” are computed on all template and complement reads together (rather than treating the two read types separately).

**Supplementary Table 3D:** Top 10 2D lengths with mean quality score (Q) across all 3 runs.

| Rank | Length | Q | Run |
| --- | --- | --- | --- |
| 1 | 102935 | 8.74 | A |
| 2 | 96237 | 9.61 | A |
| 3 | 91062 | 9.93 | B |
| 4 | 86797 | 9.01 | C |
| 5 | 84898 | 8.87 | A |
| 6 | 84716 | 8.28 | B |
| 7 | 84477 | 7.75 | A |
| 8 | 83697 | 9.34 | B |
| 9 | 82373 | 9.73 | A |
| 10 | 78658 | 8.24 | A |

**Supplementary Table 3E:** Top 10 1D reads where mean quality Q≥3

| Rank | Length | Q | Run | Read Type |
| --- | --- | --- | --- | --- |
| 1 | 210931 | 3.37 | A | Template |
| 2 | 202293 | 3.53 | A | Template |
| 3 | 199014 | 3.40 | A | Template |
| 4 | 170334 | 3.87 | A | Complement |
| 5 | 159563 | 4.30 | A | Complement |
| 6 | 159397 | 3.92 | A | Template |
| 7 | 148912 | 3.70 | A | Template |
| 8 | 143763 | 4.17 | B | Template |
| 9 | 139864 | 4.28 | C | Template |
| 10 | 132439 | 4.20 | C | Complement |

**Supplementary Table 4A:**
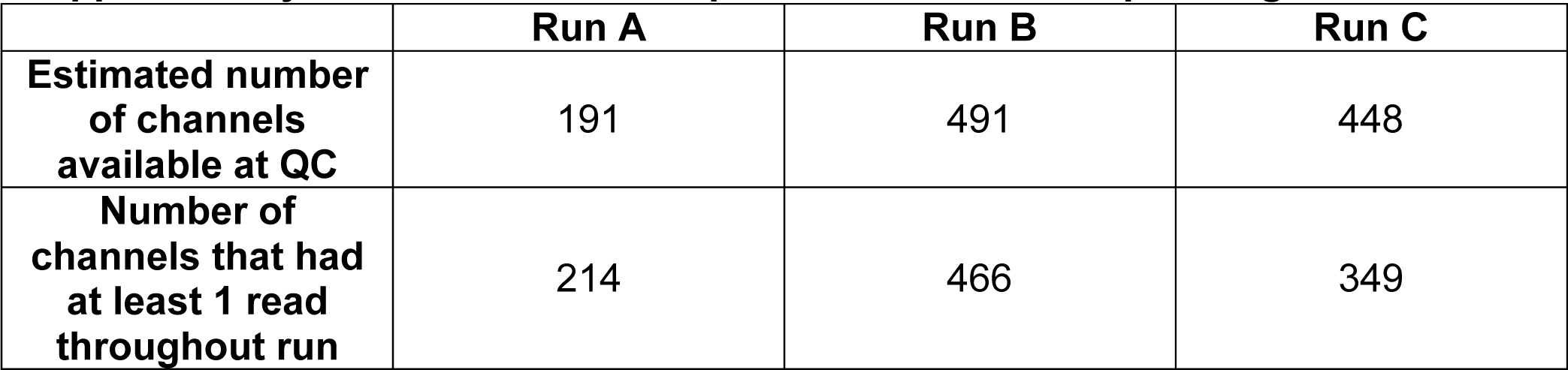
Number of pores available for sequencing.

**Supplementary Table 4B:** Schedule and recipes for adding sequencing mixes to flow cells during MinION sequencing.

|  | Run A |  |  |  | Run B |  |  |  | Run C |  |  |  |
| --- | --- | --- | --- | --- | --- | --- | --- | --- | --- | --- | --- | --- |
|  | PSM | EP | Fuel | HAP | PSM | EP | Fuel | HAP | PSM | EP | Fuel | HAP |
| **SM1** | 6 | 140 | 4 | - | 3 | 144 | 3 | - | 6 | 140 | 4 | - |
| **SM2** | 6 | 140 | 4 | 10 | 4 | 142 | 3 | 1 | 6 | 140 | 4 | 1 |
| **SM3** | 3 | 143 | 4 | 6 | 4 | 143 | 3 | 8 | 3 | 144 | 3 | 6 |
| **SM4** | 6 | 140 | 4 | 8 | 4 | 174 | 4 | 15 | 2 | 145 | 3 | 12 |
| **SM5** | 3 | 143 | 4 | 13 | 3 | 144 | 3 | 5.5 | 3 | 143 | 4 | 5 |
| **SM6** | R | 143 | 4 | 5 | 4 | 142 | 4 | 4 | R | 145 | 3 | 7 |
| **SM7** | - | - | - | - | R | 144 | 4 | 10 | - | - | - | - |

HAP = Hours After Previous (i.e. time after adding previous SM)
PSM = Pre-Sequencing Mix
SM = Sequencing Mix
EP = EP Buffer
Fuel = "Fuel Mix" provided by ONT.
R = Remaining PSM
Sequencing Mix (SM) was prepared fresh immediately before adding to the flow cell at each time point.
All volumes reported for PSM, EP, and Fuel are in μl.

**Supplementary Table 5:** Filtering.

|  | **Run A** | **Run B** | **Run C** |
| --- | --- | --- | --- |
| **Number event files** | 27667 | 82040 | 12536 |
| **Number files filtered: no template found (NTF)** | 10306<br>(37.3%) | 6820<br>(8.31%) | 1997<br>(15.9%) |
| **Number files filtered: too few events (TFE)** | 7414<br>(26.8%) | 10878<br>(13.26%) | 1583<br>(12.6%) |
| **Number of files filtered: too many events (TME)** | 9<br>(0.03%) | 46<br>(0.056%) | 14<br>(0.11%) |
| **Number of files filtered: time error (TE)** | 2<br>(0.007%) | 14<br>(0.017%) | 0<br>(0 %) |
| **Number of files with <100,000 events with time error**<br>**(percent of all files with <100,000 events)** | 0<br>(0%) | 0<br>(0%) | 0<br>(0%) |
| **Number of files with 100,000 to 1,000,000 events with**<br>**time error (percent of all files with 100,000 to 1,000,000**<br>**events)** | 2<br>(1.57%) | 14<br>(5.51%) | 0<br>(0%) |
| **Number of TME files (>1,000,000 events) that also had**<br>**time errors** | 5 of 9<br>(55.6%) | 30 of 46<br>(65.2%) | 4 of 14<br>(28.6%) |
| **Number of TME files (>1,000,000 events) that did NOT**<br>**have time errors** | 4 of 9<br>(44.4%) | 16 of 46<br>(34.8%) | 10 of 14<br>(71.4%) |
| **Number of TME files (>1,000,000 events) without time**<br>**errors that did NOT have lead adapter profile** | 0 of 4<br>(0%) | 6 of 16<br>(37.5%) | 5 of 10<br>(50%) |
| **Number of TME files (>1,000,000 events) without time**<br>**errors that had lead adapter profile** | 4 of 4<br>(100%) | 10 of 16<br>(62.5) | 5 of 10<br>(50%) |
| **Number/percent of TME files (>1,000,000 events) that**<br>**passed time-error and lead adapter filtering** | 4 of 9<br>(44.4%) | 10 of 46<br>(21.7%) | 5 of 14<br>(35.7%) |

**Supplementary Table 6A:** Are there more “0 moves” near G4 motifs than sites selected at random?

|  |  | Null 1 | Null 2 | Null 3 | Null 4 |
| --- | --- | --- | --- | --- | --- |
| Run A | Sign | 7.554388e-26 | 4.832767e-38 | 1.193051e-137 | 5.480738e-122 |
|  | Signed Rank | 3.090888e-27 | 2.244648e-41 | 4.612996e-131 | 2.224541e-124 |
| Run B | Sign | 0 | 3.482397e-315 | 0 | 0 |
|  | Signed Rank | 0 | 0 | 0 | 0 |
| Run C | Sign | 1.29939e-41 | 1.698042e-36 | 6.549949e-48 | 3.301627e-55 |
|  | Signed Rank | 2.194097e-75 | 1.639146e-65 | 2.938907e-50 | 2.956933e-50 |
| p-value product | Sign | 0 | 0 | 0 | 0 |
|  | Signed Rank | 0 | 0 | 0 | 0 |

0 indicates that $p < 1e-324$ .
G4s and stays:
G4s paired with Nulls
Null 1 = randomly selected positions on template and complement reads without G4 motifs
Null 2 = randomly selected positions on all template and complement reads
Null 3 = randomly selected positions on the same template or complement read containing the G4 motif
Null 4 = randomly selected positions on the same template or complement read containing the G4 motif excluding positions that overlap the G4 motif
Note: 7.95%, 7.84%, and 7.24% of Template and Complement reads contain G4 motif in Run A, Run B, and Run C, respectively.

**Supplementary Table 6B:** Do G4 motifs on complement strand associate with more “0 moves” than G4 motifs on template?

|  | Rank Sum |
| --- | --- |
| Run A | 1.033993e-30 |
| Run B | 3.952016e-26 |
| Run C | 0.004113 |
| p-value product | 1.680719e-58 |

**Supplementary Table 6C:** Do G4 motifs with >4 tracts associate with more “0 moves”?

|  | Rank Sum |
| --- | --- |
| Run A | 0.002023 |
| Run B | 2.139787e-92 |
| Run C | 4.449e-14 |
| p-value product | 1.925878e-108 |

**Supplementary Table 6D:**
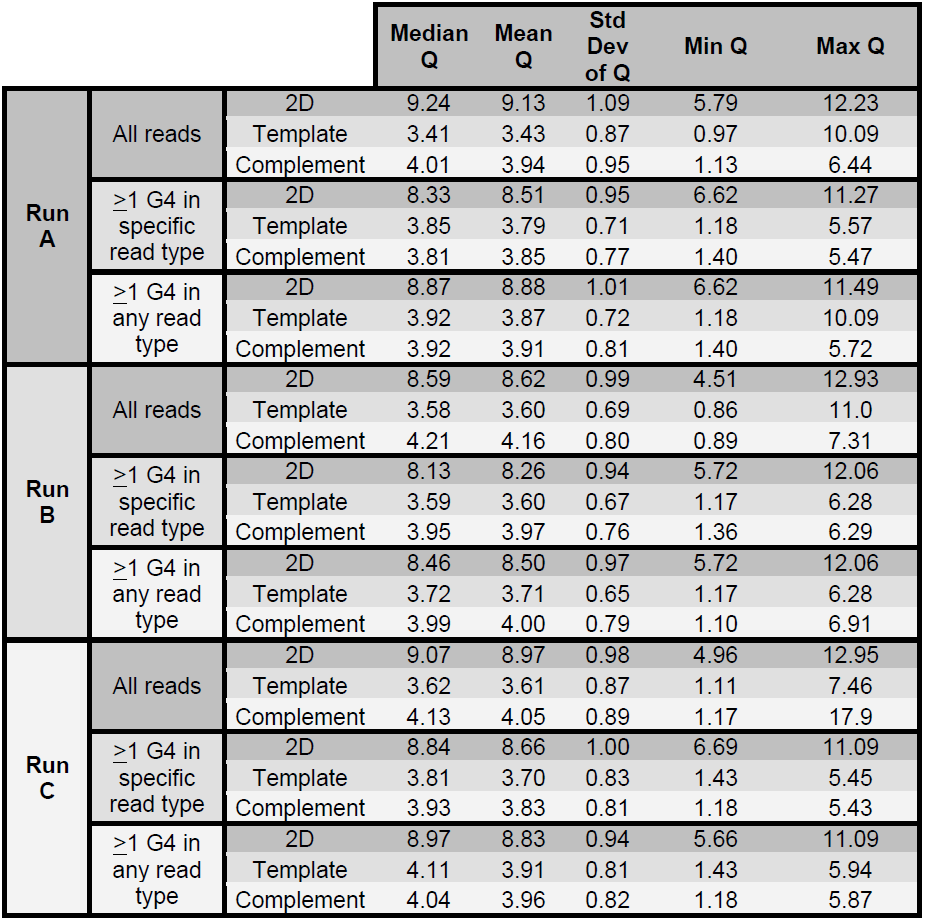
Q score distributions for all reads, specific read types with ≥1 G4 motif, any read type with ≥ 1 G4 motif.

## REFERENCES

1. Check Hayden, E. Pint-sized DNA sequencer impresses first users. Nature 521, 15–16 (2015).

2. Kilianski, A. et al. Bacterial and viral identification and differentiation by amplicon sequencing on the MinION nanopore sequencer. Gigascience 4, 12 (2015).

3. Loman, N. J. & Watson, M. Successful test launch for nanopore sequencing. Nat. Methods 12, 303–304 (2015).

4. Quick, J. et al. Rapid draft sequencing and real-time nanopore sequencing in a hospital outbreak of Salmonella. Genome Biol. 16, 114 (2015).

5. Mikheyev, A. S. & Tin, M. M. Y. A first look at the Oxford Nanopore MinION sequencer. Mol. Ecol. Resour. 14, 1097–1102 (2014).

6. Quick, J., Quinlan, A. R. & Loman, N. J. A reference bacterial genome dataset generated on the MinION(TM) portable single-molecule nanopore sequencer. Gigascience 3, 22 (2014).

7. Goodwin, S. et al. Oxford Nanopore Sequencing and de novo Assembly of a Eukaryotic Genome. *bioRxiv* (Cold Spring Harbor Labs Journals, 2015). doi:10.1101/013490

8. Ashton, P. M. et al. MinION nanopore sequencing identifies the position and structure of a bacterial antibiotic resistance island. Nat. Biotechnol. 33, 296–300 (2014).

9. Jain, M. et al. Improved data analysis for the MinION nanopore sequencer. Nat. Methods 12, 351–356 (2015).

10. Ammar, R., Paton, T. A., Torti, D., Shlien, A. & Bader, G. D. Long read nanopore sequencing for detection of HLA and CYP2D6 variants and haplotypes. F1000Research 4, 17 (2015).

11. Madoui, M.-A. et al. Genome assembly using Nanopore-guided long and error-free DNA reads. BMC Genomics 16, 327 (2015).

12. Warren, R. L., Vandervalk, B. P., Jones, S. J. & Birol, I. LINKS: Scaffolding genome assemblies with kilobase-long nanopore reads. bioRxiv (Cold Spring Harbor Labs Journals, 2015). doi:10.1101/016519

13. Bolisetty, M., Rajadinakaran, G. & Graveley, B. Determining Exon Connectivity in Complex mRNAs by Nanopore Sequencing. bioRxiv (Cold Spring Harbor Labs Journals, 2015). doi:10.1101/019752

14. Chiu, C. Y. et al. Rapid metagenomic identification of viral pathogens in clinical samples by real-time nanopore sequencing analysis. bioRxiv (Cold Spring Harbor Labs Journals, 2015). doi:10.1101/020420

15. Cao, M. D. et al. Real-time strain typing and analysis of antibiotic resistance potential using Nanopore MinION sequencing. bioRxiv (Cold Spring Harbor Labs Journals, 2015). doi:10.1101/019356

16. Szalay, T. & Golovchenko, J. A. A de novo DNA Sequencing and Variant Calling Algorithm for Nanopores. bioRxiv (Cold Spring Harbor Labs Journals, 2015). doi:10.1101/019448

17. Sovic, I. et al. Fast and sensitive mapping of error-prone nanopore sequencing reads with GraphMap. bioRxiv (Cold Spring Harbor Labs Journals, 2015). doi:10.1101/020719

18. Loman, N. J., Quick, J. & Simpson, J. T. A complete bacterial genome assembled de novo using only nanopore sequencing data. bioRxiv (Cold Spring Harbor Labs Journals, 2015). doi:10.1101/015552

19. Check Hayden, E. Nanopore genome sequencer makes its debut. Nature (2012). doi:10.1038/nature.2012.10051

20. Check Hayden, E. Data from pocket-sized genome sequencer unveiled. Nature (2014). doi:10.1038/nature.2014.14724

21. Huppert, J. L. Structure, location and interactions of G-quadruplexes. FEBS J.277, 3452–8 (2010).

## Supplementary References

1. Team, R. C. R: A language and environment for statistical computing. (R Foundation for Statistical Computing, 2013).

2. Loman, N. J. & Quinlan, A. R. Poretools: a toolkit for analyzing nanopore sequence data. Bioinformatics btu555– (2014). doi:10.1093/bioinformatics/btu555

3. Huppert, J. L. Structure, location and interactions of G-quadruplexes. FEBS J. 277, 3452–8 (2010).

4. Huppert, J. L. & Balasubramanian, S. Prevalence of quadruplexes in the human genome. Nucleic Acids Res. 33, 2908–16 (2005).

5. Quinlan, A. R. & Hall, I. M. BEDTools: a flexible suite of utilities for comparing genomic features. Bioinformatics 26, 841–2 (2010).

